# Reticular Adhesion Formation is Mediated by Flat Clathrin Lattices and Opposed by Fibrillar Adhesions

**DOI:** 10.1101/2022.05.30.494022

**Authors:** Laura Hakanpää, Amr Abouelezz, An-Sofie Lenaerts, Seyda Culfa, Michael Algie, Jenny Bärlund, Pekka Katajisto, Harvey T. McMahon, Leonardo Almeida-Souza

**Affiliations:** Helsinki Institute of Life Science, HiLIFE, University of Helsinki, Helsinki, Finland; Institute of Biotechnology, University of Helsinki, Helsinki, Finland; Faculty of Biological and Environmental Sciences, University of Helsinki, Helsinki, Finland; Department of Cell and Molecular Biology, Karolinska Institutet, Stockholm, Sweden; MRC Laboratory of Molecular Biology, Cambridge, UK

## Abstract

Reticular adhesions (RAs) consist of integrin αvβ5 and harbor flat clathrin lattices (FCLs), long-lasting structures with similar molecular composition to clathrin mediated endocytosis (CME) carriers. Why FCLs and RAs colocalize is not known. Here, we show that FCLs assemble RAs in a process controlled by fibronectin (FN) and its receptor, integrin αvβ5. We observed that cells on FN-rich matrices displayed fewer FCLs and RAs. CME machinery inhibition abolished RAs and live-cell imaging showed that RA establishment requires FCL co-assembly. The inhibitory activity of FN was mediated by the activation of integrin α5β1 at Tensin1-positive fibrillar adhesions. Conventionally, endocytosis disassembles cellular adhesions by internalization of their components. Our results present a novel paradigm in the relationship between these two processes by showing that endocytic proteins can actively function in the assembly of cell adhesions. Furthermore, we show this novel adhesion assembly mechanism is coupled to cell migration via a unique crosstalk between cell matrix adhesions.

## Introduction

Integrins are nonenzymatic dimeric transmembrane receptors which recognize extracellular matrix (ECM) components. These mechanosensory proteins govern cell adhesion to the ECM maintaining correct tissue development and function, with elaborate connections to cellular homeostasis and disease (Kanchanawong and Calderwood, 2023). Ligand availability, and biochemical and physical properties of the ECM, determine integrin activation status, integrin clustering, and, ultimately, the formation of cellular adhesion structures (Kechagia et al., 2019).

Cells can form a variety of integrin-based adhesions. Small Integrin clusters engaged to the ECM, called nascent adhesions, form on the cell periphery and establish their connection to the actin cytoskeleton via adaptor proteins such as Talin. A balancing act of traction forces and signaling molecules determine whether nascent adhesions mature into the larger and molecularly more complex focal adhesions (FAs) (Wehrle-Haller, 2012). In migrating cells and in the presence of the ECM component fibronectin (FN), FAs can serve as platforms for the formation of fibrillar adhesions (FB), where FN-bound α5β1 integrins “slide” from FAs to form FN fibrils (Georgiadou and Ivaska, 2017). In common to all these types of cell adhesions, their disassembly is mediated by the removal of integrin molecules from adhesion sites via endocytosis (Kechagia et al., 2019).

Recently, a novel type of integrin-based cell adhesion was discovered (Lock et al., 2018). Called reticular adhesions (RAs), these structures contain integrin αvβ5, lack the typical markers for the other adhesion types, such as Talin1 or Paxillin, and are not connected to actin stress fibers. RAs can occupy a significant portion of the substrate-facing surface of cells in culture and can significantly outlast FAs. Their physiological function is, however, not clear.

Intriguingly, RAs colocalize with large, persistent forms of clathrin structures at the cell membrane called Flat Clathrin Lattices (FCLs) (also referred to as clathrin plaques) (Grove et al., 2014). The structure containing FCLs and RAs is called clathrin-containing adhesion complexes (CCAC) (Lock et al., 2019; Zuidema et al., 2020). FCLs were previously considered as stalled endocytic events of the Clathrin-Mediated Endocytosis (CME) pathway. However, recent studies have changed this view and support the idea that FCLs are signaling platforms (Leyton-Puig et al., 2017; Grove et al., 2014; Alfonzo-Méndez et al., 2022). In vivo, FCLs localize to adhesive structures between bone and osteoclasts (Akisaka et al., 2008) and are required for the organization of sarcomeres (Vassilopoulos et al., 2014).

The functional relationship between FCLs and RAs is not clear. A confounding factor in this relationship lies in the fact that, although FCLs always localize to RAs, the opposite is not true. RAs can occur as large structures with FCLs covering only a fraction of their area. Moreover, integrin αvβ5 can localize to both RAs and FAs. Although details on the factors mediating integrin αvβ5 localization to FCLs are becoming clearer (Zuidema 2018 and 2022), why these structures co-exist, what their function is and how cells control their formation remain a mystery.

In this study, we show that FCLs are required for the establishment of RAs. Moreover, we found that a FN-rich ECM acts as an inhibitor of FCL-mediated RA formation. This inhibitory role of FN is mediated by the activation of integrin α5 β1 localized at fibrillar adhesions. Furthermore, we show that the transition from a static to a migratory state is mirrored by the disappearance of FCLs and RAs.

## Results

### Fibronectin inhibits the formation of FCLs

While studying CME dynamics, we serendipitously observed that cells on fibronectin (FN) appear to display fewer FCLs when compared to cells plated on non-coated glass dishes. To confirm if ECM proteins in general can influence CME, we assessed the effect of several major ECM components as well as non-ECM coatings and non-coated surfaces on the amount of FCLs. For that, dishes were coated for 16-24 h after which cells were let to attach for 16-20 h in serum-containing medium before imaging. For quantifications, we established the metric *FCL proportion*, which defines the average fraction of FCLs per frame among all clathrin coated structures detected in a 5-minute movie (see methods for details). These experiments were performed using U2Os cells with an endogenously GFP-tagged alpha adaptin 2 sigma subunit (AP2S1, hereafter referred to simply as AP2). AP2 is a widely used CME marker which faithfully mirrors clathrin dynamics (Almeida-Souza et al., 2018; Ehrlich et al., 2004; Rappoport and Simon, 2008). We used endogenously tagged cell lines throughout this study as the expression level of the AP2 complex was shown to modulate the amount of FCLs (Dambournet et al., 2018).

U2Os cells on non-coated dishes presented typical and abundant FCLs (i.e. bright, long-lived AP2-GFP marked structures) (Figure 1A, B, S1A and Supplementary video 1), similar to what has been found in many cell lines (Zuidema et al., 2022; Moulay et al., 2020; Sochacki et al., 2021; Saffarian et al., 2009). Similarly, U2Os cells plated on dishes coated with the non-ECM proteins bovine serum albumin (BSA) and poly-L-lysine (PLL) also presented high FCL proportions (Figure 1B, S1A and Supplementary video 1). Out of the major ECM proteins tested, FN, collagen IV (Col IV) and laminin-111 (LN111) reduced FCL proportion significantly. The integrin αvβ5 ligand vitronectin (VTN) did not increase or decrease the FCL proportion when compared to non-coated dishes (Figure 1B, S1A) (see discussion). Similarly, and in line with a recent study (Baschieri et al., 2018), collagen I (Col I) did not reduce FCLs (Figure 1 B, S 1 A and Supplementary video 1). Different concentrations of FN used for coating (10 or 20μg/ml) did not show significant differences (Figure 1B).

**Figure 1.**
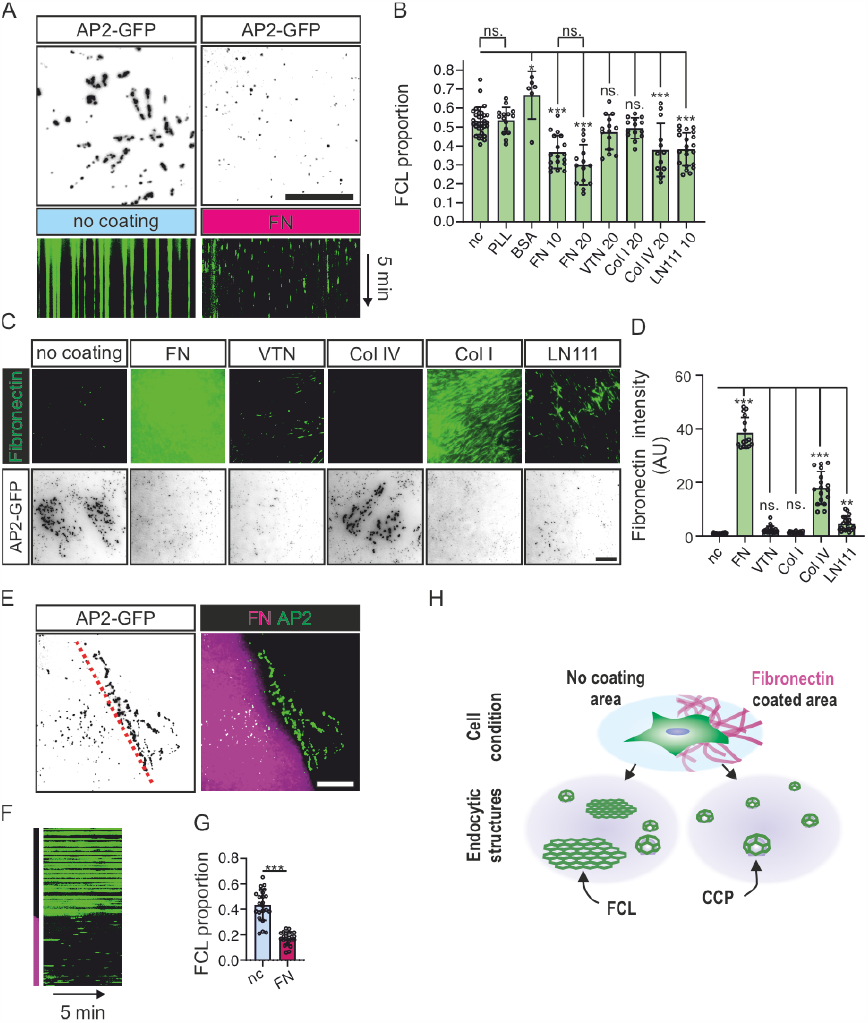
Fibronectin inhibits FCL formation in a local manner. (A) U2Os-AP2-GFP cells plated on non-coated or FN-coated dishes were imaged using TIRF microscopy, at 1 s intervals for 5 min. Images represent 15 s time projections, and kymographs represent the 5 min time-lapse videos. (B) FCL proportions of U2Os-AP2-GFP cells plated on PLL, BSA, FN, VTN, Col I, Col IV, LN111-coated or non-coated dishes and imaged using TIRF microscopy, at 1 s intervals for 5 min. N (videos): non-coated (nc)=32, PLL 3ug/ml=6, BSA 20ug/ml=15, FN 10 μg/ml=18, FN 20 μg/ml=14, VTN 20 μg/ml=14, Col I 20 μg/ml = 14, Col IV 20 μg/ml=14, LN111 10 μg/ml=22. Videos were collected from four independent experiments, except for PLL, from two experiments. One-way ANOVA with Tukey’s multiple comparison test. F(8, 183)=19.11, P<0.0001. (C) U2Os-AP2-GFP cells plated on FN, VTN, Col I, Col IV, LN111 (all 10 μg/ml)-coated or non-coated dishes were and stained for FN. Representative TIRF images. (D) Quantification of fibronectin fluorescent intensity from samples in C. N (images): FN=15 images, VTN/Col I/LN111/non-coated=21 images; Col IV=17 images. Results were obtained from one representative experiment. Similar results were observed in four individual experiments. F(5, 110)=338.5, P<0.0001. (E) U2Os-AP2-GFP cells plated on FN/glass patterned imaging dishes and imaged with TIRF microscopy at 1 s intervals for 5 min. Representative images are 15 s time projections. (F) Representative 5 min kymograph of time-lapse videos in E. (G) Quantification of FCL proportions in E. n=23 videos, collected from 4 individual experiments. Two-tailed Student’s t-test, P<0.001. (H) A schematic illustration of results shown in this figure. Data are the mean ± SD, ns. non-significant p-value; * p-value < 0.05; p-value < 0.01; *** p-value < 0.001. Scalebars 10 μm.

Recently, it was described that SCC-9 cells produce more FN when plated on Col IV or LN111 (Lu et al., 2020). To probe if this is also the case for our cells, we stained FN from U2Os cells plated directly onto non-coated dishes, or plated on FN, VTN, Col IV, Col I or LN111. While U2Os cells produce little FN overnight, cells plated on Col IV produced a striking amount of FN, which assembled into elongated fibrils (Figure 1C and 1D). LN111 coating also induced FN production, but less strikingly than Col IV. Col I and VTN coating were unable to stimulate FN production (Figure 1C, D). These results suggest that FN is the main ECM component inhibiting FCL formation.

For many cell lines, it is common to find considerable variability in the amount of FCLs in culture. We thus decided to test if this variability is due to differential FN production within the culture. Confirming this hypothesis (and bearing in mind that U2Os secrete FN modestly, see below), we found that cells plated on non-coated dishes displaying fewer FCLs were predominantly lying on top of a FN-rich region of the culture (Figure S1B).

Next, we asked if the reduction in FCL proportions observed in FN-coated samples are a cell-wide effect or specific to cellular regions in direct contact with the extracellular substrate. For that, we used patterned dishes containing FN-coated regions interspersed with uncoated regions, where single U2Os-AP2-GFP cells could adhere simultaneously to both a FN- and a non-coated region. In line with a contact-dependent effect, low FCL proportions were observed in cellular regions in contact with FN whereas FCL proportion was high in cellular regions contacting non-coated surface (Figure 1E-G and Supplementary video 2).

The abundance of an alternative splice variant of the clathrin heavy chain containing exon 31 was recently shown to increase the frequency of clathrin plaques in myotubes (Moulay et al., 2020). We thus tested if the effects we observe are also due to changes in clathrin splicing, but found no difference when comparing cells plated on non-coated or FN-coated dishes (Figure S1C). Thus, these results show that FN is a potent inhibitor of FCLs. Moreover, FN inhibits FCLs in a contact-dependent manner locally within a single cell (Figure 1H).

### Fibronectin inhibits the formation of RAs in a similar manner as FCLs

As discussed in the introduction, FCLs localize to RAs. To check how ECM composition affects these structures, U2Os AP2-GFP cells plated on FN, VTN, Col IV, Col I or LN111 and stained with the RA component integrin αvβ5 and-to be able to distinguish integrin αvβ5 on RAs or FAs-were also stained with an FA marker (phosphorylated paxillin, p-PAX Y118). Cells plated overnight without coating formed abundant RAs (Figure 2A, B). On FN-coated dishes, big RAs were largely absent but small “dot-like” nascent RAs were present in a few cells. Similarly, on Col IV and LN111 coatings (which stimulated FN production) (Figure 1C), cells formed significantly fewer RAs than on non-coated dishes (Figure 2A, B). Coating with VTN, the ligand for integrin αvβ5 present at RAs and FAs alike (Figure S1C) (Lock et al), did not result in more RAs (Figure 2A, B). Different coatings also changed the total amount of integrin αvβ5 on the bottom surface of cells (Figure S1D). However, they did not follow a clear relationship with the amount of RAs. To quantify differences in RA amounts in cells we developed a metric called *RA coverage*, which measures the fraction of the area of the cell covered by integrin αvβ5 signal (excluding FAs). RA coverage serves as a good metric to distinguish between large and nascent RAs and, crucially, it shows a clear correspondence between RA content and both FCL proportion and FN abundance in the ECM (see figures Figure 1B, 1D and 2B).

**Figure 2.**
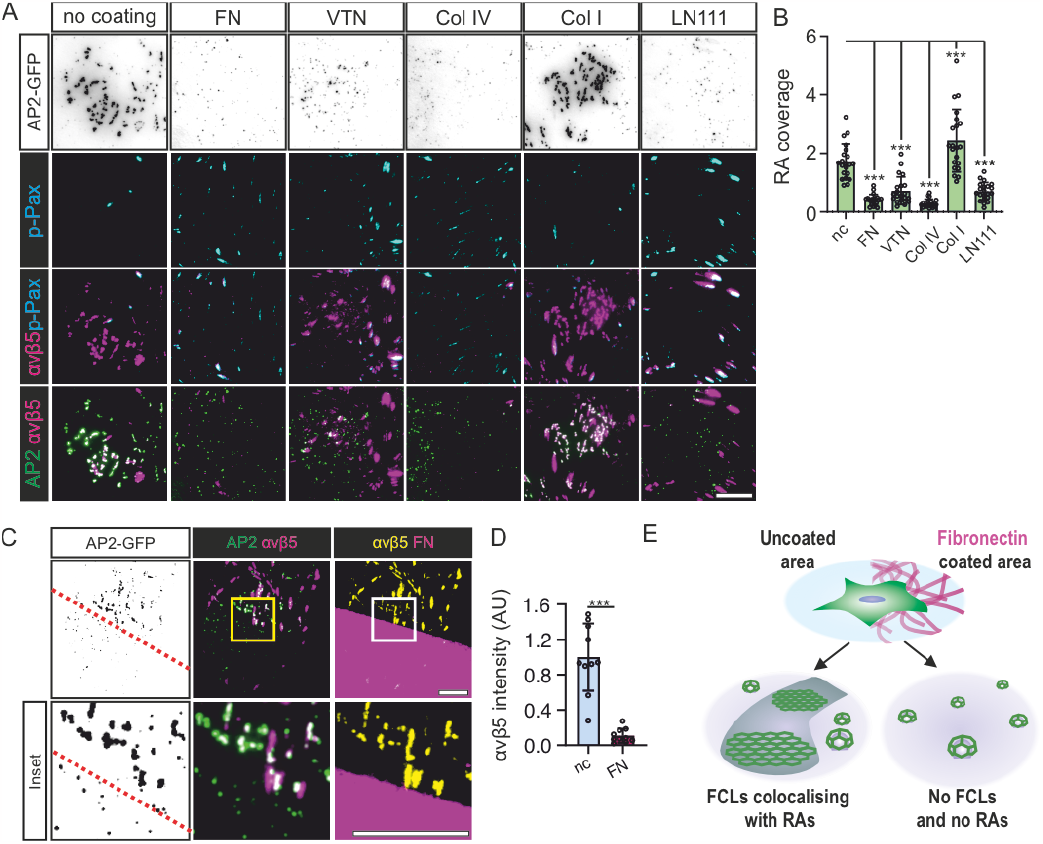
Fibronectin inhibits RA formation in a local manner. (A) U2Os-AP2-GFP cells plated on FN, VTN, Col I, Col IV, LN111 (all 10 μg/ml)-coated or non-coated dishes, stained for p-PAX and integrin αvβ. Representative TIRF images. (B) Analysis of RA coverage from samples in A. N=21 from two independent experiments. F(5, 120)=48.05, P<0.0001. (C) U2Os-AP2-GFP cells were plated on FN/glass patterned imaging dishes over night, stained for integrin αvβ5 and p-Pax, and imaged using TIRF microscopy, representative TIRF images. (D) Quantification of integrin αvβ5 fluorescent intensity on FN- and glass-side of the pattern, n=10 from one representative experiment. Similar results were observed in five similar experiments. Two-tailed Student’s t-test, P<0.01. (E) A schematic illustration of results shown in this figure. Data are the mean ± SD, ns. non-significant p-value; *** p-value < 0.001. Scalebars 10 μm, insets 5 μm.

Next, we used our substrate patterning strategy to check if the local FN effects on FCLs were also similar for RAs. Strikingly, cells plated on patterned FN revealed that RAs, akin to FCLs, were completely inhibited on cellular regions in contact with FN. Cellular regions in contact with non-coated surfaces displayed many FCLs colocalizing to RAs while regions in contact with FN presented no RAs or FCLs (Figure 2C). Interestingly, in these patterned substrates, most of the integrin αvβ5 signal segregated to non-coated regions forming typical RAs (Figure 2C,D). This contrasts with cells plated in fully coated dishes (Figure 2A), where integrin αvβ5 can be seen in both RAs and FAs. Hence, the inhibitory effects of FN on FCLs affects RAs in a similar manner (Figure 2E).

### The effect of fibronectin on FCLs and RAs is clear in various cell lines

Next, we checked if the effects we see in U2Os cells are also true for other cell lines. To avoid problems of overexpression, we endogenously tagged AP2 with either Halo tag or GFP in various human cell lines: HeLa (epithelial, cervical carcinoma), MCF7 (epithelial, breast cancer), HDF (dermal fibroblast, noncancerous), Caco2 (epithelial, colon carcinoma) and hMEC (Human mammary epithelial cells). These cell lines presented a large variation in the amount of FCLs and the morphology of RAs. Importantly, these cells could be divided into two groups in terms of endogenous FN secretion, and this division clearly correlated with the amount of FCLs and RAs (Figure 3A, B). U2Os, HeLa and MCF7 composed the group of low FN-secretion cells. U2Os form large RAs on non-coated dishes, whereas HeLa formed multiple dot-like nascent RAs (which colocalized with FCLs) with bigger RAs found more seldom (Figure 3A). MCF7 cells formed many FCLs and large RAs covering almost the entire cell area (Figure 3A). None of the high FN-secreting cell lines (HDF, Caco2 and hMEC) formed large RAs (Figure 3A, B). In these high FN-producing cells, small FCL/RA dots were often found in areas with less deposited FN (Figure 3A).

**Figure 3.**
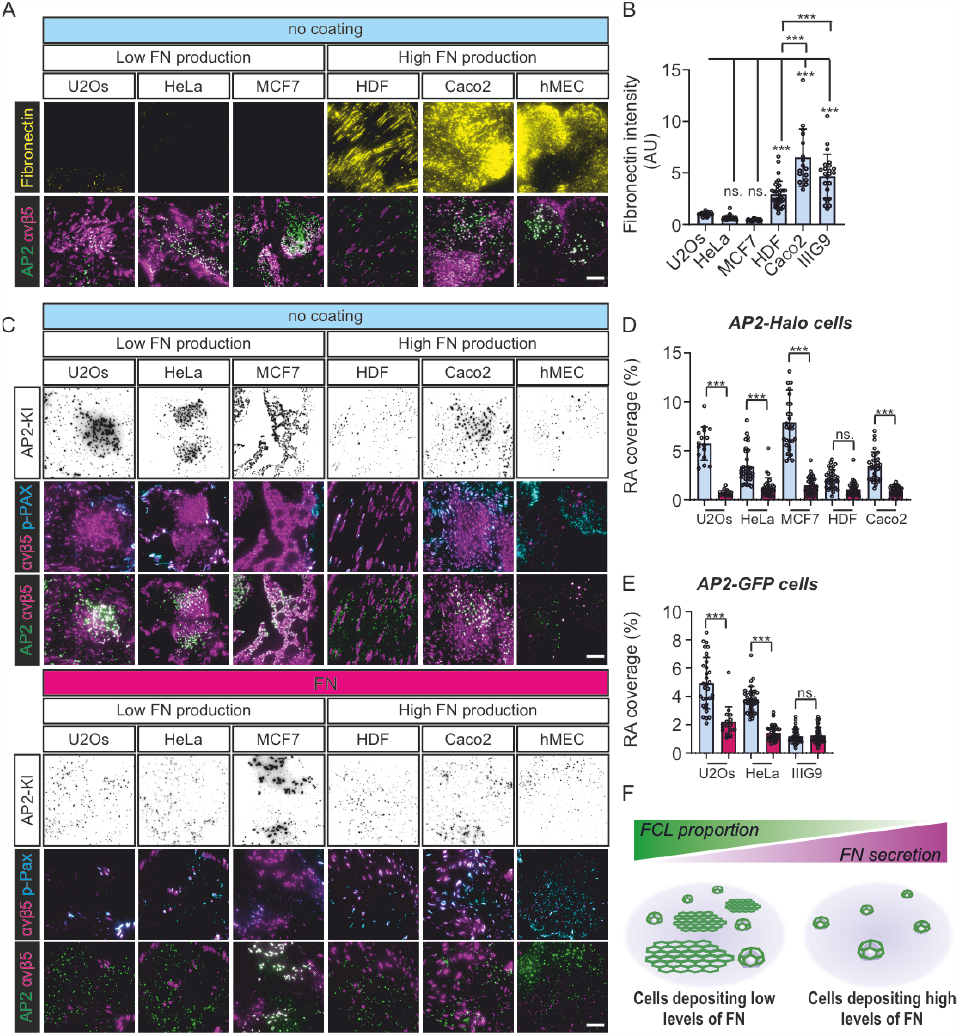
FCLs and RAs presence correlates with FN production in multiple cells lines. (A-E) The following knock-in cell lines were used in this figure: U2Os-AP2-GFP, HeLa-AP2-GFP, hMEC-AP2-GFP, U2Os-AP2-halo, HeLa-AP2-halo, MCF7-AP2-halo, HDF-AP2-halo, and Caco2-AP2-halo. (A) Cell lines indicated were plated to non-coated dishes, allowed to settle overnight and stained for integrin αvβ5 and FN. Representative TIRF images. (B)Analysis of FN fluorescent intensity from samples in A. N (images): U2Os=21, HeLa=20, MCF7=23, HDF=33, Caco2=16, hMEC=23, from two independent experiments. F(5, 130)=56.16, P<0.0001. (C) Cells were plated on FN-coated or non-coated dishes, allowed to settle overnight, and stained for integrin αvβ5 and p-Pax. Representative TIRF images. (D) Analysis of RA coverage from samples in C. Cell lines with an AP2 halo tag, n (images): U2Os (nc/FN)=20, HeLa (nc)=33, HeLa (FN)=29, MCF7 (nc)=29, MCF7 (FN)=32, HDF (nc/FN) 32, Caco2 (nc/FN)=32. Data were obtained from two individual experiments and similar results were observed in four individual experiments. F(9, 273)=64.96, P<0.0001. (E) Analysis of RA coverage from samples in C. Cell lines with an AP2 GFP tag, n (images): U2Os (nc)=31, U2Os (FN)=18, HeLa (nc)=35, HeLa (FN)=38, hMEC (nc/ FN)=38. Data were obtained from two individual experiments and similar results were observed in four individual experiments. F(5, 189)=89.97, P<0.0001. (F) A schematic illustration of results shown in this figure. Data are the mean ± SD, ns. non-significant p-value; *** p-value < 0.001. Scalebars 10 μm.

We next evaluated the response of these cell lines to FN pre-coating. In low FN-producing cell lines (U2Os, HeLa, and MCF7), RA coverage dropped significantly (Figure 3C-E). Among the high FN-production cells lines, only Caco2 reduced its RA coverage on FN-coated dishes (Figure 3C,D). As expected, HDF and hMEC, which had low RA coverage without coating, did not show a significant response to FN coating (Figure 3C, E).

For all experiments so far, we used media supplemented with serum, which is known to contain ECM components, including FN. Given that our cells are left to attach overnight in this media, it would be reasonable to expect that the FN present in serum would coat the dishes and completely mask our results. To test why this does not seem to happen (Figures 1C and 3A), we compared the amount of FN deposited on the glass surface in different conditions: dishes were coated for 24 h with 10, 5 and 1μg/ml of FN (diluted in PBS), 100% fetal bovine serum (FBS), media with 10% FBS and PBS as a control. After coating, U2Os cells were plated and left to attach for 16 h before being fixed and stained for FN. Surprisingly, our results revealed that very little FN was deposited on glass in dishes “coated” with full media or pure FBS (Figure S1F-G). These results are in line with similar experiments performed 30 years ago (Steele et al., 1992). We hypothesize that this phenomenon occurs due to the high concentrations of BSA in serum (40 mg/ml) which rapidly saturates the surface of culture dishes, thereby acting as a blocking agent for the binding of serum FN.

Taken together, the presence of FN controls the formation of RAs and FCLs in a very similar manner and in different cells lines (Figure 3F), suggesting a common and general-mechanism for the establishment of these structures.

### The CME machinery is essential for RA formation

Next, we set out to dissect the relationship between the formation of FCLs and RAs. It has been shown that integrin αvβ5 is required for the establishment of FCLs (Baschieri et al., 2018; Zuidema et al., 2018). We confirmed this observation by silencing integrin β5 from U2Os AP2-GFP cells plated on non-coated dishes and, indeed, they displayed a significantly lower FCL proportion compared to control cells (Figures S2A, B). Further, while integrin β5-silenced cells were unable to form RAs, they did form FAs (Figure S2C, D). The dependency of integrin β5 on FCL formation was further confirmed using Cilengitide, the inhibitor for integrin αvβ5 (Desgrosellier and Cheresh, 2010), as the treatment led to a rapid disassembly of FCLs and RAs (Figures S2E-F).

While all FCLs colocalize to RAs, FCL-free areas of larger RAs are rather common (e.g. Figures 2A, 3A, 3C and S2D), which may give the impression that FCLs are formed on pre-existing RAs. Nevertheless, the fact that both structures are inhibited independently by FN suggests a deeper relationship and led us to ask if RAs can exist without the CME machinery. To answer this question, we quantified the RA coverage in U2Os-AP2-GFP cells silenced for the clathrin adaptor AP2 complex subunits alpha 1 (AP2A1) or sigma 1 (AP2S1) in cells plated on non-coated dishes, a condition where we observe large RAs. Consistent with an important role played by the CME machinery in RA formation, AP2A1-or AP2S1-silenced cells (easily recognizable as cells with little to no AP2-GFP signal), did not display RAs. Instead, integrin αvβ5 localized to FAs (Figure 4A, B).

**Figure 4.**
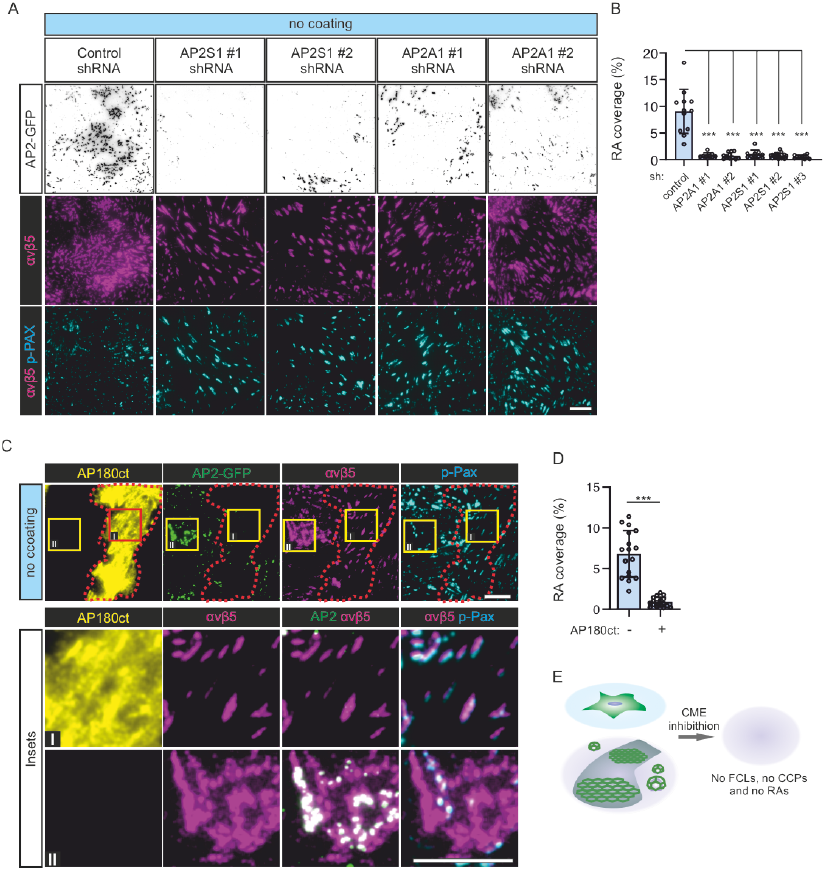
Inhibition of CME prevents RA formation. (A) U2Os-AP2-GFP cells silenced for AP2 subunits alpha 1 (AP2A1), sigma 1 (AP2S1) or control (scrambled shRNA) were plated on non-coated dishes and stained for integrin αvβ5 and p-Pax. Representative TIRF images. (B) Analysis of RA coverage from samples in A. N (images): control=12, shAP2A1#1=8, shAPA1#2=11, shAPS1#1=10, shAPS1#2= 19, shAPS1#3=10. Data were obtained from two individual experiments and similar results were observed in five independent experiments. One-way ANOVA with Tukey’s multiple comparison, F(5, 64)=43.11, P<0.001. (C) U2Os-AP2-GFP cells overexpressing Ap180ct were plated on non-coated dishes and stained for integrin αvβ5 and p-Pax. Representative TIRF images. (D) Analysis of RA coverage from samples in C, n=17 from one representative experiment and similar results were observed in four individual experiments. Two-tailed Student’s t-test, P<0.0001. (E) A schematic illustration of results shown in this figure. Data are the mean ± SD,*** p-value < 0.001. Scalebars 10 μm, insets 5 μm.

To confirm these results, we expressed the AP180 C-terminal fragment (AP180ct), which acts as a strong dominant negative of CME (Ford et al., 2001). AP180ct-positive U2Os-AP2-GFP cells plated on non-coated dishes displayed low AP2 signal at the membrane and, akin to AP2-silenced cells, RAs were largely absent with integrin αvβ5 localized to FAs, whereas AP180ct-negative cells displayed typical FCLs and RAs (Figure 4 C, D). Thus, the CME machinery is required for the formation of RAs (Figure 4E).

Next, we set out to visualize the dynamics of AP2 during RA formation. For that, we generated a double U2Os knock-in cell line-AP2-GFP and Integrin β5 (ITGB5)-mScarlet. RAs are remarkably stable structures (Lock et al., 2018) and their de novo formation is rare, making it rather difficult to capture such events. To minimize this challenge, we optimized the conditions for Cilengitide treatment to disassemble RAs followed by a washout, when RAs could start reforming (Figures S3A, B). Using these washout conditions, we were able to capture events showing that the formation and growth of ITGB5-postive structures are accompanied by the formation of FCLs (Figures 5A, S3C and S3D and Supplementary videos 3 and 4). Also, in events where we could not clearly detect the extension of a mature RA (reminiscent of the dot-like structures we see in many cells), we noticed that the establishment of an FCL was typically accompanied by an increase in ITGB5 fluorescence (Figures 5B, S3C and S3D and Supplementary video 3). Importantly, ITGB5-postive structures, which did not colocalise with an FCL, rapidly disappeared. In many cases, this disappearance was preceded by bona fide CME events (short lived AP2-GFP signals), likely representing CME-mediated ITGB5-cluster disassembly (Figures 5B and S3D).

**Figure 5.**
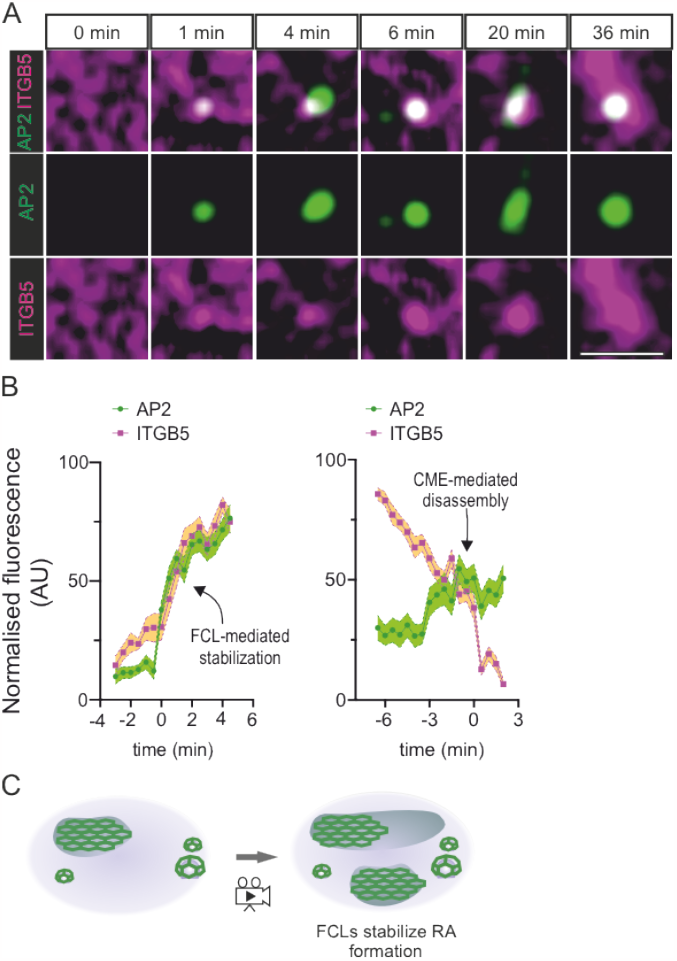
RAs are formed at FCLs. (A) U2Os-AP2-ITGB5-mScarlet cells plated on glass were treated with Cilengitide (10 μM) for 15 minutes, washed twice with fresh medium and imaged live using TIRF microscopy at 2 frames per minute. Representative frames showing the growth of an RA from an FCL. (B)Analysis of AP2 and integrin αvβ5 (ITGB5) intensity over time. Left: FCL-mediated RA assembly (n=16 events). Right: ITGB5 clusters not stabilized by FCLs disassemble and are removed from the membrane by CME-mediated disassembly (n=24 events). Events have been collected from four 1 h long time-lapse acquisitions. Mean ± SEM. (C) A schematic illustration of results shown in this figure. Scalebar 5 μm.

Taken together, our results show that the relationship between FCLs and RAs is beyond a simple colocalization. In fact, our data reveals a strict co-dependency, where FCLs are required for the stabilization and growth of integrin αvβ5 clusters thereby establishing RAs (Figure 5C).

### *The inhibitory effect of fibronectin on FCL and RA formation is mediated by integrin* α5β1

To understand the mechanism controlling the co-assembly of FCLs and RAs, we turned our attention back to FN. While integrin αvβ5 binds to VTN at FAs and RAs, the major FN receptor is integrin α5β1 (Humphries et al., 2006). First, we acutely interfered with integrin β1 binding to FN using the function-blocking antibody mab13. U2Os-AP2-GFP cells seeded on FN-coated dishes were treated with mab13 and monitored for the acute formation of FCLs and RAs. Over the time course of 45 min, mab13 induced the relocalization of integrin αvβ5 from FAs to small, newly formed RAs (Figure 6A, B). Further supporting the role of FCLs in RA assembly, these newly formed RAs completely colocalized with FCLs (bright AP2 signals) (Figure 6A, C). A similar experiment followed by live-cell imaging confirmed these results and showed a gradual increase in FCL proportions after mab13 treatment (Figure 6D). Mab13 treatment had no effect on existing FCLs and RAs formed on non-coated dishes (data not shown).

**Figure 6.**
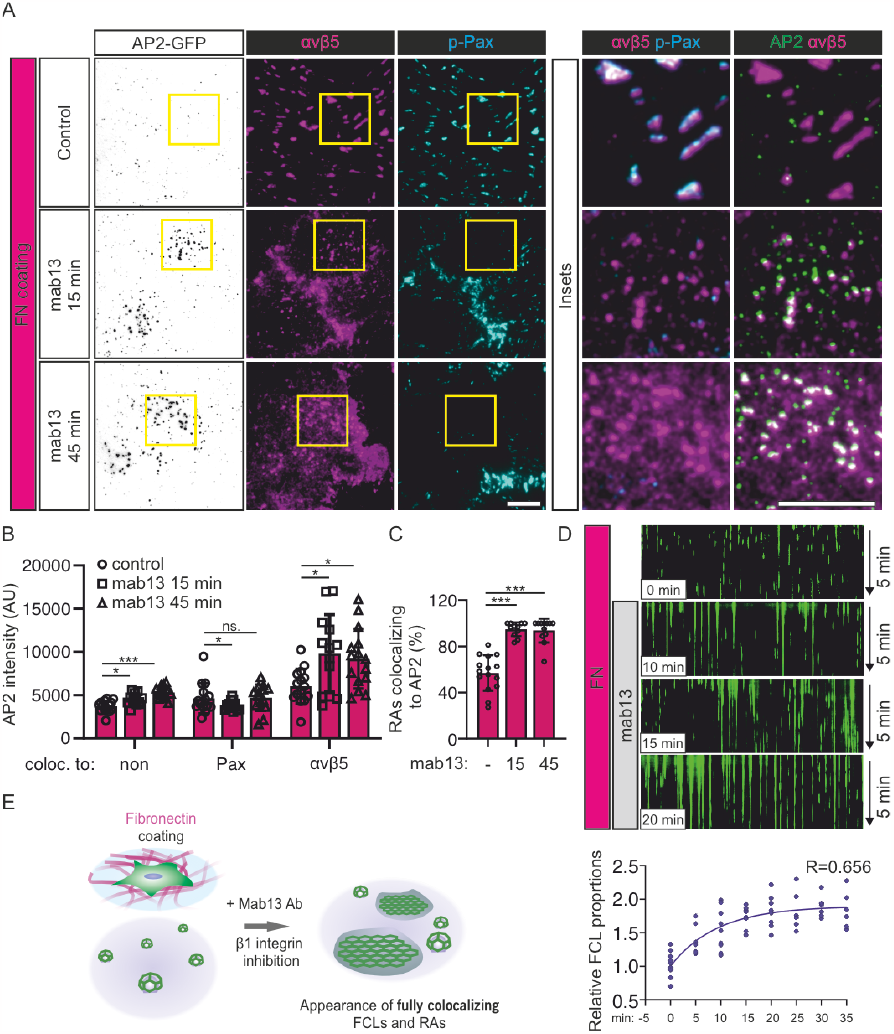
Integrin β1 blocking stimulates FCL and RA formation. (A) U2Os-AP2-GFP cells plated on FN were treated with integrin β1 blocking antibody mab13 (0.3 μg/ml) for 15 and 45 min (or vehicle for 45 min=control) and stained for integrin αvβ5 and p-Pax. Representative TIRF images. (B) Analysis of AP2 signal colocalising with no markers (non), p-Pax or integrin αvβ5 over time. N(images): control=18, mab13 15 min=13, mab13 45 min=15, from one representative experiment. Similar results were observed in four independent experiments. One-way ANOVA with Tukey’s multiple comparison. F(2, 119)=46.68, P<0.0001. (C) Analysis of fraction of integrin αvβ5 colocalising with AP2 over time from samples in A. N (images): control=15, mab13 15 min= 11, mab13 45 min=12, from one representative experiment. Similar results were observed in four independent experiments. One-way ANOVA with Tukey’s multiple comparison. F(2, 35)=45.81, P<0.0001. (D) U2Os-AP2-GFP cells plated on FN were treated with mab13 (0.3 μg/ml) and imaged using TIRF microscopy. 5 min time-lapses with 1 s intervals starting at 0 min (no mab13) and every 5 min after mab13 addition, until 35 min, were acquired. N (videos): 0 min=11, 5min=11, 10 min=9, 15 min=8, 20 min= 8, 25 min=6, 30 min=7, 35 min=6. Videos were acquired from two independent experiments. Similar results were observed in five individual experiments. One-way ANOVA with Tukey’s multiple comparison. F(7, 29)=8.893, P<0.0001. (E) A schematic illustration of results shown in this figure. Data are the mean ± SD, ns. non-significant p-value; * p-value<0.05, *** p-value < 0.001. Scalebars 10 μm, insets 5 μm.

In line with these results, integrin β1 silencing in U2Os-AP2-GFP cells plated on FN resulted in a high FCL proportion and large, prominent RAs (Figure 7A-D and S4A-C). Despite the striking increase of integrin αvβ5 on the bottom surface of silenced cells, this increase was not reflected in expression levels, indicating that the stimulation of RA formation leads to a change in the trafficking of this integrin dimer (Figure S4C).

**Figure 7.**
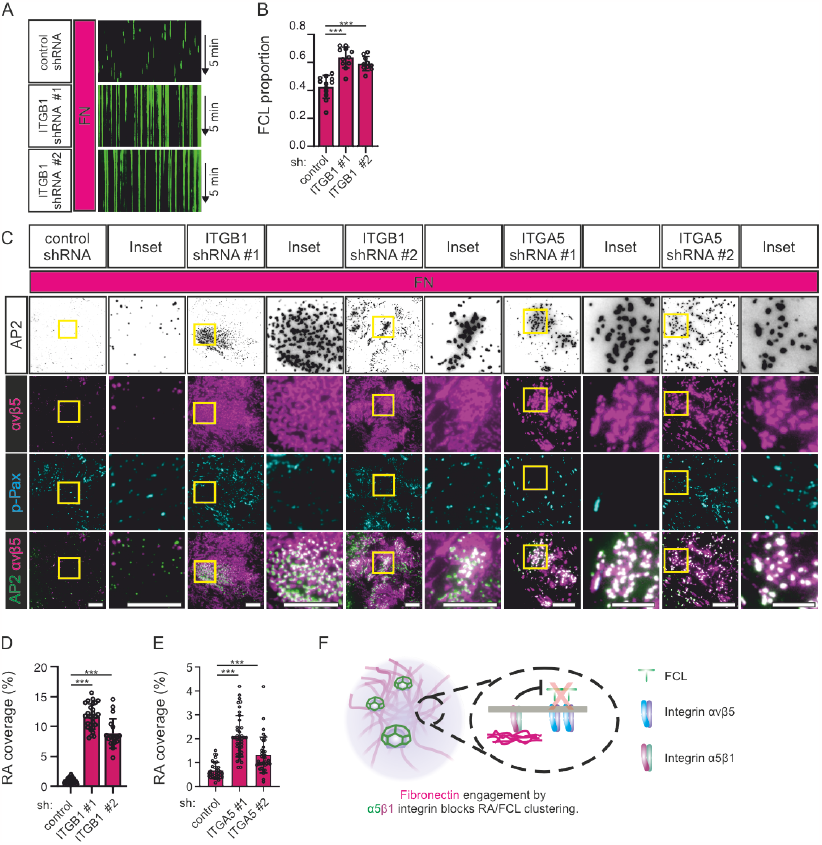
Depletion of integrin β1 promotes FCL and RA formation. (A) U2Os-AP2-GFP cells silenced for integrin β1 with two different shRNAs (shITGB1 #1, #2) or control shRNA were plated on FN-coated dishes and stained for p-Pax and imaged using TIRF microscopy at 1 s intervals for 5 min. Representative 5 min kymographs. (B) Analysis of FCL proportions from samples in A. N (videos): shScr=12, shITGB1 #1=11, shITGB1 #2=10, from three individual experiments. Tukey’s multiple comparison, F(2, 30)=27.81, P<0.001.(C) U2Os-AP2-GFP cells silenced for integrin β1 with two different shRNAs (shITGB #1, #2), or integrin a5 with two different shRNAs (shITGA5 #1, #2), or control shRNA, were plated on FN-coated dishes and stained for integrin αvβ5 and p-Pax. Representative TIRF images. (D) Analysis of RA coverage from integrin β1 silenced samples in C. N (images): control shRNA=30, shITGB1 #1=27, shITGB1 #2=18, from two independent experiments, similar results were observed in four independent experiments. One-way ANOVA with Tukey’s multiple comparison, F(2, 72)=276.2, P<0.0001. (E) Analysis of RA coverage from integrin a5 silenced samples in E. N (images): control shRNA=33, shITGA5 #1=40, shITGA5 #2=37, from three individual experiments. One-way ANOVA with Tukey’s multiple comparison, F(2, 107)=44.46, P<0.0001. (F) A schematic illustration of results shown in this figure. Data are the mean ± SD, *** p-value < 0.001. Scalebars 10 μm, insets 5 μm.

A significant increase in RAs was also seen in cells silenced for integrin a5, the alpha subunit which pairs with integrin β1 for FN binding (Figure 7C, E and S4D, E). Taken together, these results show that the inhibitory activity of FN on RAs and FCLs occurs via the activation of integrin α5β1 (Figure 7F).

### *Activation of integrin* α5β1 *at fibrillar adhesions controls RA and FCL formation*

When bound to FN, Integrin α5β1 can slide centripetally on the cell membrane, translocating from FAs to form elongated structures called fibrillar adhesions (FB). This movement generates long FN fibrils in a process called FN fibrillogenesis and is mediated by the cytoskeleton scaffolding protein Tensin1 (Pankov et al., 2000). To determine which type of adhesion structure active integrin α5β1 localizes to under our experimental conditions, we plated U2Os-AP2-GFP cells on FN and non-coated dishes and stained them with an active integrin β1-specific antibody (12G10) and Tensin1 or p-Pax to mark FBs or FAs, respectively. The staining revealed that in the FN-coated dishes, active integrin β1 was colocalizing to FBs (Tensin1) (Figure 8A). As expected, active integrin β1 and Tensin1-positive adhesions were largely absent in non-coated dishes (Figure 8A).

**Figure 8.**
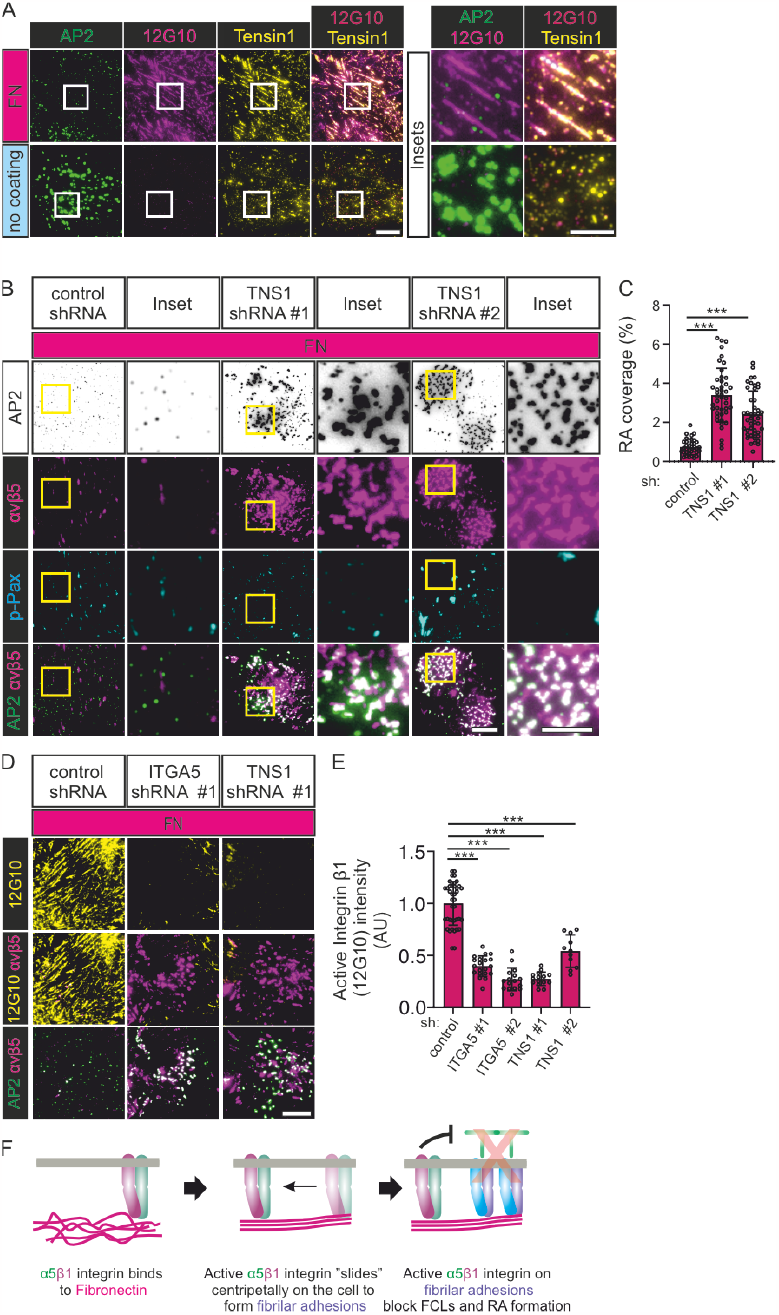
Active integrin α5β1 at fibrillar adhesions inhibit FCL and RA formation. (A) U2Os-AP2-halo cells were plated to FN-coated or non-coated dishes and stained for Tensin1 and active integrin β1 12G10. Representative TIRF images. (B) U2Os-AP2-GFP-ITGB5-mScarlet cells silenced for Tensin1 with two different shRNAs (shTNS1 #1, #2) or control shRNA, were plated on FN-coated dishes and stained for p-Pax. Representative TIRF images. (C) Analysis of RA coverage from samples in B. N (images): shScr control=35, shTNS1 #1/#2=44, from three independent experiments. One-way ANOVA with Tukey’s multiple comparison, F(2, 120)=56.26, P<0.0001. (D) U2Os-AP2-GFP-ITGB5-mScarlet cells silenced for Tensin1 with two different shRNAs (shTNS1 #1, #2) or control shRNA were plated on FN-coated dishes and stained for active integrin β1 (12G10 antibody). Representative TIRF images. (E) Analysis of 12G10 fluorescent intensity from samples in D. N (images): shScr=20, shTNS1 #1=15, shTNS1 #2=11, from one representative image. Similar results were observed in three individual experiments. One-way ANOVA with Tukey’s multiple comparison, F(2, 43)=87.81, P<0.0001. (F) A schematic illustration of results shown in this figure. Data are the mean ± SD,*** p-value < 0.001. Scalebars 10 μm, insets 5 μm.

Next, to determine which active integrin β1 pool is more important for the inhibition of FCLs and RAs, we silenced FAs and FB components on U2Os-AP2-GFP cells and plated them on FN. In accordance with the higher accumulation of active integrin β1 in FBs, silencing of Tensin1 led to a marked increase of RAs and FCLs accompanied by a reduction in the presence of active integrin β1 on the membrane (evidence by 12G10 antibody staining) (Figure 8B, C and S5A-C). Silencing of the FA component Talin-1 also led to increased RAs and FCLs and a reduction of active integrin β1 on the membrane (Figure S5D-F). Given the strong phenotype on Tensin1 knockdown, this result was expected as FAs are precursors of FBs.

FB formation indicates activated and migratory cell phenotypes. Indeed, active sliding of integrin α5β1 and Tensin1 bound to FN along central actin stress fibers increases traction forces (Georgiadou and Ivaska, 2017; Pankov et al., 2000) and is required in cell migration during development and cancer metastasis (Efthymiou et al., 2020; Schwarzbauer and DeSimone, 2011). If the extension of active integrin α5β1 into FBs is indeed required for the inhibition of FCLs and RAs, we hypothesized that physical confinement of cells — which inhibits cell migration — would also inhibit the sliding of FB from FAs. In turn, the absence of integrin α5β1 in FBs would favor FCLs and RAs, even if cells were plated on an FN-rich matrix. To test this possibility, we turned to single cell micropatterns. In contrast to the patterned coatings we used in Figures 1 and 2, these micropatterns do not allow cells to attach outside the defined areas on a coverslip. Given the small size of these areas (1100 μm^2^), cells are laterally confined. U2Os-AP2-GFP cells were plated on slides with arrow- and H-shaped micropatterns either precoated with FN or not and stained for integrin αvβ5 and p-PAX and imaged to measure RA coverage. In addition, to measure integrin β1 activation, U2Os-AP2-GFP-ITGB5-mScarlet cells were plated similarly and stained for active integrin β1. Supporting our hypothesis, we could detect clear FCL and RAs in FN-coated micropatterns (Figure 9A, B, S5K). On arrows, FCLs and RAs developed on the shaft of the pattern, rather than the arched area. In the H-patterns, FCLs and RA developed all over the pattern. Cells on non-coated patterns made large FCLs and RAs often extending throughout the pattern (Figure 9A, B). Crucially, the RA coverage was not significantly different between coated or non-coated patterns (Figure 9B). As expected, staining with active integrin β1 (12G10) showed a clear difference in signal between FN-coated and non-coated patterns (Figure 9C, S5L). Importantly, further supporting a need for FB formation to inhibit FCLs and RAs, 12G10 signal was not organized as elongated, central FBs but rather confined at the cell periphery (Figure S5K). Thus, the inhibitory role of FN on FCLs and RA formation occurs primarily via the activation of integrin α5β1 on Tensin1-positive fibrillar adhesions.

**Figure 9.**
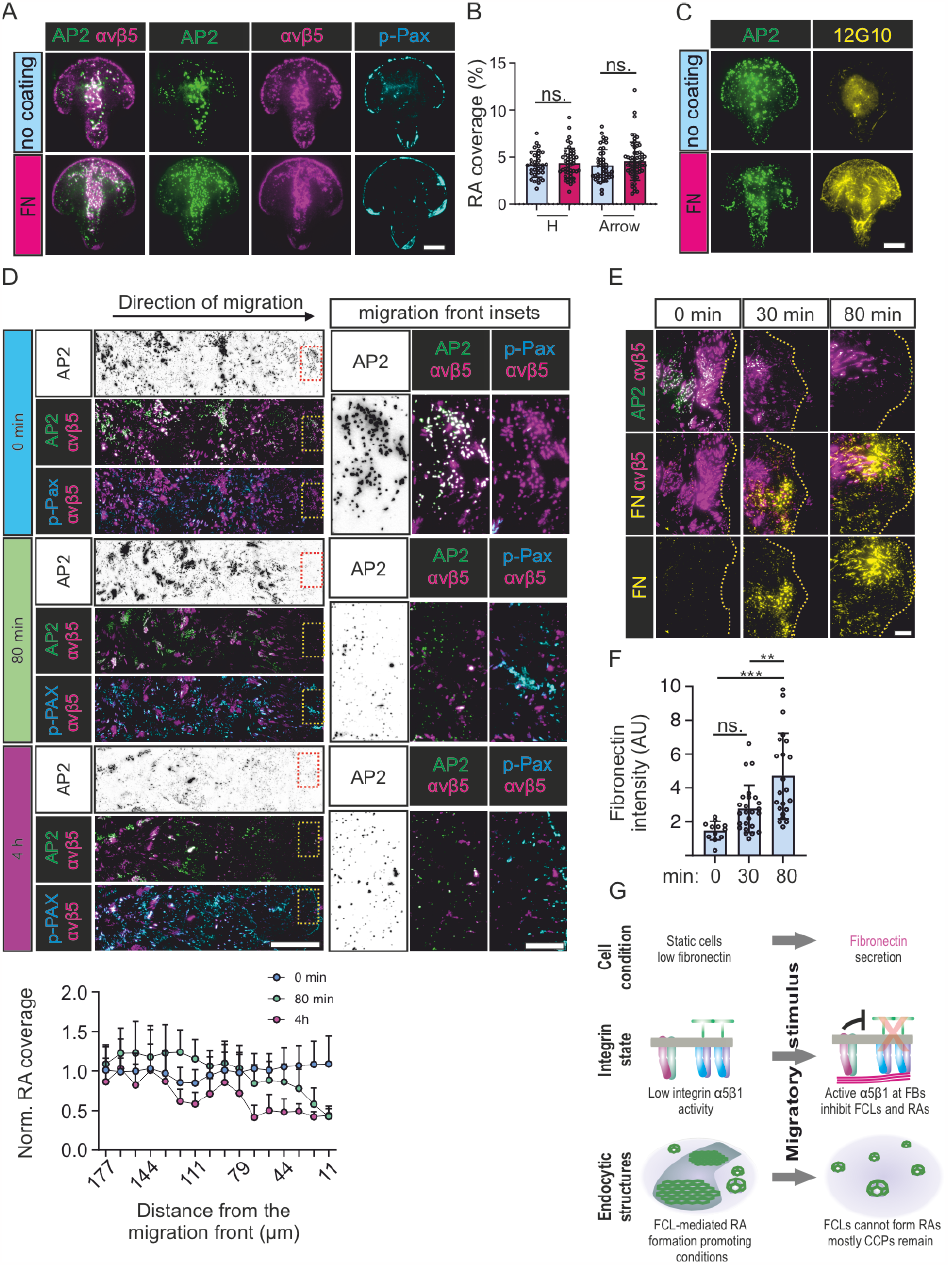
RA disassembly is coupled to cell migration. (A) U2Os-AP2-GFP cells were grown on FN-precoated or non-coated micropatterns (1100 mm2) and stained for integrin αvβ5 and p-Pax. Representative TIRF images. See figure S5K for the same staining of H-shaped patterns. (B) Analysis of RA coverage from samples in A. N (images): Arrow non-coated=28, Arrow FN=27, H non-coated=25, H FN=20, from one representative image. One-way ANOVA with Tukey’s multiple comparison, F(2, 79)=38.09, P<0.0001. (C)U2Os-AP2-GFP-ITGB5-mScarlet cells were grown on FN-coated or non-coated micropatterns (1100 mm2) and stained for active integrin β1 12G10. Representative TIRF images. See figure S5L for the same staining with H-shaped patterns. (D) top: U2Os-AP2-GFP-ITGB5-mScarlet cells were grown on non-coated dishes for 2 d, wounded, and let to migrate for 80 min or 4 h in fresh complete medium, and stained for p-Pax. Representative TIRF images (stitched tile of 5 side-by-side fields of view). The wound is exactly at the right edge of the images. (D) Bottom: Analysis of normalized RA coverage from tiles in D. RA coverage was calculated in 11 μm wide sliding windows from the wound edge. N (tiles): 0 min=48, 80 min=36, 4 h=15. One-way ANOVA with Tukey’s multiple comparison, F(2, 79)=38.09, P<0.0001. (E) U2Os-AP2-GFP-ITGB5-mScarlet cells were grown on non-coated dishes for 2 d, wounded, and let to migrate for 30 min or 80 min in fresh complete medium, and stained for FN. Representative TIRF images from the migrating front (indicated as yellow lines). (F) Analysis of FN fluorescent intensity from samples in E. N (images): 0 min=10, 30 min=25, 80 min=22. One-way ANOVA with Tukey’s multiple comparison, F(2, 56)=13.77, P<0.0001. (G) A schematic illustration of results shown in this figure. Data are the mean ± SD,** p-value < 0.01, *** p-value < 0.001. Scalebars 10 μm, except in D 50 μm, and inset 10 μm.

### The disassembly of FCL/RA is coupled to cell migration

As physical restriction favored FCLs and RAs, we wondered if inducing migration will have the opposite effect. To test this hypothesis, we monitored FCLs and RAs in a classic wound healing assay. U2Os-AP2-GFP-ITGB5-mScarlet cells were plated on non-coated dishes and allowed to grow to full confluency for 2 days. Cultures were then wounded and cells were allowed to migrate. At 0 minutes (i.e. just after wounding), FCLs and RAs were abundant and equally distributed at the edge and away from the wound (Figure 9D). Within 80 minutes, the cells at the migration front had lost most of their FCLs and RAs, while cells further away from the edge maintained their FCLs and RAs (Figure 9D). At 4 hours, as the migratory front grew larger, the loss of FCL and RAs also extended away from the wound (Figure 9D). In full accordance with the results we presented above, the disappearance of FCLs and RAs was preceded by the increase in FN secretion by the cells at the edge of the wound (Figure 9E, F). Together, these results place the resolution of FCLs and RAs as an intrinsic part of the cascade of events triggering cell migration (Figure 9G).

## Discussion

The extracellular environment is a key regulator of cellular physiology with integrins playing a key role translating the chemical composition of the extracellular milieu into intracellular signals. Among various mechanisms controlling integrin function, integrin trafficking via endocytosis and exocytosis plays a major role (Moreno-Layseca et al., 2019). Thus far, the relationship between integrin-based matrix adhesions and endocytosis has been considered primarily antagonistic, with endocytosis playing a role in the disassembly of said adhesive structures (Ezratty et al., 2009; Moreno-Layseca et al., 2019). Here we provide evidence, for the first time, of a constructive relationship between the endocytic machinery and cellular adhesions, where the CME machinery, in the form of FCLs, is key for the formation of integrin αvβ5 RAs. Moreover, we show that FCL-mediated αvβ5 RA formation is counteracted by the activation of a distinct integrin heterodimer, α5β1, in distinct adhesion structures, FBs, revealing an interesting mechanism of inter-adhesion crosstalk.

Our results support the idea that FCLs and RAs are two sides of the same structure (Lock et al., 2019; Zuidema et al., 2020). Previous studies have demonstrated the importance of integrin αvβ5 at RAs in the formation of FCLs (Zuidema et al., 2022, 2018; Lock et al., 2019). Here, we show that this relationship is also crucial in the other direction, with FCLs being required for the formation of integrin αvβ5 RAs. Therefore, we believe the previously suggested term clathrin containing adhesion complexes (or CCAC for short) is a more appropriate terminology to refer to these structures.

### The mechanism of FCL-mediated RA formation

We observed that FCL-mediated RA formation events are rare, which led us to use non-physiological conditions-a Cilengitide washout experiment-to detect them. Therefore, the physiological trigger leading to the formation of FCLs and establishment of RAs remains to be understood. VTN, the ligand for integrin αvβ5 could be considered a good candidate. However, as we show in figure 2E and as reported by others (Zuidema et al., 2022), integrin αvβ5 binds VTN equally on FAs and RAs. While it is clear that the presence of VTN is important as an extracellular tether for the formation of integrin αvβ5 adhesions (Zuidema et al., 2022, 2018; Lock et al., 2018), the switch between these adhesion types is likely an inside-out mechanism. We did not detect an increase in integrin αvβ5 RAs on VTN-coated dishes (Figure 1B). This could seem counterintuitive, but VTN-which was initially called “serum spreading factor” (Hayman et al., 1983)-is readily secreted by cells during attachment. Therefore, we advise caution when making conclusions based on the results of non-coated and VTN-coated dishes on the role of this ECM component on RA coverage. Further work is necessary to shed light on this issue.

Recent evidence showed that EGFR activation led to the enlargement of FCLs in an integrin β5 phosphorylation-dependent manner (Alfonzo-Méndez et al., 2022), pointing to a possible mechanism for the initial co-assembly of FCLs and RAs. This possibility is further reinforced by the fact that the relationship between growth factor receptors and integrins has been established in multiple contexts (Ivaska and Heino, 2011).

Another key unknown aspect of FCL-mediated RA formation concerns how these structures can be molecularly differentiated from canonical endocytic events. The connection between integrin αvβ5 located in RAs and FCLs occurs primarily via the endocytic adaptors ARH and NUMB (Zuidema et al., 2018). Importantly, these adaptors also participate in integrin endocytosis (Ezratty et al., 2009; Nishimura and Kaibuchi, 2007), suggesting that other mechanisms may be required to define the identity of FCLs.

Recently, a correlation was found between the presence of clathrin plaques and an alternatively spliced isoform of clathrin containing exon 31 in myotubes (Moulay et al., 2020). We did not detect any changes in clathrin splicing in our experimental system, which was not surprising given the effects we see are contact-dependent and could not be explained by transcriptional changes. In addition, we cannot ensure that the clathrin plaques detected in myotubes are equivalent to the FCLs we observe here. Nonetheless, it is possible that the abundance of the exon 31-positive clathrin isoform works as a dial that changes the probability, speed or efficiency by which cells form FCLs.

### Another unusual function for the clathrin machinery

In addition to its endocytic function, the clathrin machinery has been shown to participate in other processes. For example, clathrin helps to stabilize the mitotic spindle by binding to microtubules (Royle, 2012) and, during E. coli infection, the CME machinery is co-opted to form a clathrin-based actin-rich adhesive structure for the bacteria called a pedestal (Veiga et al., 2007). Furthermore, a clathrin/AP2 tubular lattice was recently described to envelop collagen fibers during cell migration (Elkhatib et al., 2017). The results we present here add to this list of non-endocytic functions of CME components with an important twist. FCLs can also be disassembled into individual endocytic events (Lampe et al., 2016; Tagiltsev et al., 2021; Maupin and Pollard, 1983), providing an elegant and efficient mechanism for cells to switch the same machinery from an adhesion assembly to an adhesion disassembly function.

### Inhibition of clathrin-containing complexes (CCAC) and its relationship to cell migration

In addition to defining FCLs as key factors in the establishment of CCACs, our work has also revealed many interesting aspects on the inhibition and disassembly of these structures. We show that activation of integrin α5β1 by FN and the capacity of this integrin heterodimer to slide on the plasma membrane to form fibrillar adhesions are both essential conditions for the inhibition and disassembly of CCAC. In a classical wound healing assay, we observed that as cells start to migrate they secrete FN, leading to the disappearance of CCACs. However, using laterally confined cells-which cannot form fibrillar adhesions-we observed that the mere presence of FN is not enough to inhibit CCACs. A recent study showed that high levels of activated myosin light chain (p-MLC) correlated with integrin αvβ5 localizing to FAs over RAs (Zuidema et al., 2022). Moreover, overexpression of a constitutively active RhoA mutant in a cell line with low p-MLC levels promoted integrin αvβ5 localization to FAs (Zuidema et al. 2022). As Integrin α5β1-mediated FN fibrillogenesis is required for optimal activation of the RhoA-MLC pathway, which in turn increase actin stress fiber-based migration along fibrillar adhesions (Gagné et al., 2020; Huveneers et al., 2008; Danen et al., 2002), these findings perfectly complement our data. Together, these results suggest that the disappearance of CCACs is the result of a computation of multiple cellular signals occurring during the cell migration process. Whether the disassembly of CCACs occurs actively or is a mere consequence of a non-permissive environment for the *de novo* formation of new adhesions is still unknown.

Given the fact that RAs are long-lasting cellular adhesion structures, it is tempting to hypothesize that these structures act as a “parking brake” for a cell. As the cell is triggered to migrate, this “brake” needs to be released for efficient cell movement. This process would be analogous to the loss of cell-cell contacts which happens during epithelial to mesenchymal transition (Kalluri and Weinberg, 2009), but instead of happening between cells, it would happen between the cell and the ECM. Therefore, we propose that disassembly of RAs is an intrinsic process during cells migration.

We showed that FN regulates CCAC assembly in all the cell lines we tested. However, how these *in vitro* findings will operate *in vivo* is still unknown. Even though the ECM composition in tissues is complex, the FN effect on CCAC formation is local and strictly contact dependent, which opens the possibility that, *in vivo*, tissues may use focal changes in ECM composition to control these structures.

## Author contributions

LH designed research and performed most experiments.

AA performed experiments and performed all matlab image analysis. A-SL designed and generated all knock-in cell lines.

S.C. performed and analyzed all western blots.

MA performed the experiments for the detection of alternative splicing of clathrin.

JB and PK helped with the strategy for isolation of knock-in cell lines by flow cytometry. HTM supervised the project during the initial observation.

LA-S designed research, conduct image analysis with imageJ scripts and supervised the project. LH and LA-S wrote the manuscript with input from all authors.

## Acknowledgments

We would like to thank the HILIFE light microscopy unit and the HiLIFE Flow Cytometry unit for technical assistance. We would like to thank Pekka Lappalainen, Markku Hakala and Tai Arima for the critical and kind reading of our manuscript. LA-S is supported by HiLIFE, the Academy of Finland (Research Fellow), Sigrid Juselius Foundation (Young PI grant), Finnish Diabetes Research Foundation, Magnus Ehrnrooth Foundation and Instruct-ERIC (R&D research award).

## Materials and Methods

### Cell culture and reagents

U2Os, U2Os-AP2-GFP, U2Os-AP2-halo, U2Os-AP2-GFP-ITGB5-mScarlet, HeLa-AP2-GFP, and HeLa-AP2-halo, were cultured in MEM supplemented with 10% fetal bovine serum (FBS) (Gibco) and penicillin-streptomycin (100 U/ml, Thermo Scientific). HDF-AP2-halo and Caco2-AP2-halo were cultured in DMEM supplemented with 10 % fetal bovine serum (FBS) (Gibco) and penicillin-streptomycin (100 U/ ml, Thermo Scientific). MCF7-AP2-halo was cultured in DMEM supplemented with 10% fetal bovine serum (FBS) (Gibco) and penicillin-streptomycin (100 U/ml, Thermo Scientific), 2mM glutamine, 5 μg/ml human insulin and 1μM sodium pyruvate. hMEC-AP2-GFP was cultured in MEGM complete medium (Lonza).

The following primary antibodies were used: anti-human integrin β1 clones 12G10 (Novus bio, NB100-63255), mAb13 (BD, 552828), and total integrin β1 (MAB2252, Millipore), anti-human integrin αvβ5 clone 15F11 (Millipore MAB2019Z), anti-human integrin a5 clone SNAKA51 (R&D, AF1846), anti-human Tensin1 (SAB4200283, Sigma), anti-human p-paxillin Y118 (Cell Signaling, 69363), anti-human-Talin1 (T3287, Sigma), anti-a-tubulin (sc-32293, SantaCruz Biotechnology), and anti-GAPDH (G9545, Sigma). Corresponding secondary antibodies raised against rabbit or mouse IgG were purchased from Jackson Immunoresearch.

### Coating of imaging dishes

To compare the effects of major extracellular matrix proteins on AP2 life-time, the glass coverslip areas (14 mm diameter) of imaging dishes (Mattek) were pre-coated with 10 or 20 μg/ml (300 μl in PBS), of the following ECM proteins: recombinant human fibronectin (Merck, 341631), recombinant human vitronectin (PeproTech, 140-09), collagen IV (Santa cruz, se-29010), collagen I (Sigma, C3867) and laminin-111 (Biolamina, #LN111). Coatings were incubated overnight at +37°C, except Col I was incubated in RT overnight, and LN111 was incubated in +4°C overnight. Alternatively, as non-ECM protein controls, 1% BSA (Thermo Fisher, A34785) or 0.3% poly-L-lysine (MP Biochemicals, 152690) were used to coat the dishes overnight at +37°C. Throughout this study, the standard FN coating was always performed similarly, 10 μg/ml (300 μl in PBS), overnight at +37°C.

#### FN patterning

To study local vs. global effects of FN, FN was mixed with 50 ng/ml of Alexa647-labelled BSA and used to precoat the imaging dishes overnight in +37°C. The coated surface was subsequently scratched with a needle to allow partial reappearance of non-coated surface. After scratching, the dishes were heavily rinsed with PBS. 20 000 U2Os-AP2-GFP cells were seeded on patterned imaging dishes to ensure sufficient single-cell attachment to border areas.

### Overexpression of mammalian proteins

The clathrin inhibitor AP180 c-terminal fragment (AP180ct; amino acids 516-898) cDNA, from rat origin, was described previously(Ford et al., 2001). This construct was cloned into Gateway compatible pCI vectors, containing an N-terminal monomeric EGFP using the Gateway system.

Transient transfections were carried out with PEI MAX transfection reagent (Polysciences, 24765-1) using 70% confluent U2Os cells.

### Genetic engineering of cell lines

#### Generating the U2Os-AP2-GFP and hMEC-AP2-GFP cell lines

Three guide RNA (gRNA) sequences (Integrated DNA Technologies) were designed using the Welcome Sanger Institute Genome online editing tool (https://wge.stemcell.sanger.ac.uk//https://wge.stemcell.sanger.ac.uk/). gRNAs were cloned into pSpCas9(BB)-2A-Puro (PX459) V2.0 (gift from. Feng Zhang, Addgene #62988) using BbsI sites and confirmed by Sanger sequencing.

gRNAs were transfected with the donor template for homologous recombination and the most effective gRNA (TGCTACAGTCCCTGGAGTGA), judged by the percentage of fluorescent cells by FACS, was used for single clone selection, genotyping and confirmation by microscopy.

The donor template sequence was:

GGCCAGCATCCTGGGGGGCCTCGTCTCACCCCAGGGTCTCCCCTCACACAGGTTTACACGGTCGTGGACGAGATGTTCCTGGCTGGCGAAATCCGAGAGACCAGCCAGACGAAGGTGCTGAAACAGCTGCTGATGCTACAGTCCCTGGAGGGAAGTGCATCTGGGAGCTCAGGCGCTAGTGGTTCAGCGAGCGGGGTGAGCAAGGGCGAGGAGCTGTTCACCGGGGTGGTGCCCATCCTGGTCGAGCTGGACGGCGACGTAAACGGCCACAAGTTCAGCGTGTCCGGCGAGGGCGAGGGCGATGCCACCTACGGCAAGCTGACCCTGAAGTTCATCTGCACCACCGGCAAGCTGCCCGTGCCCTGGCCCACCCTCGTGACCACCCTGACCTACGGCGTGCAGTGCTTCAGCCGCTACCCCGACCACATGAAGCAGCACGACTTCTTCAAGTCCGCCATGCCCGAAGGCTACGTCCAGGAGCGCACCATCTTCTTCAAGGACGACGGCAACTACAAGACCCGCGCCGAGGTGAAGTTCGAGGGCGACACCCTGGTGAACCGCATCGAGCTGAAGGGCATCGACTTCAAGGAGGACGGCAACATCCTGGGGCACAAGCTGGAGTACAACTACAACAGCCACAACGTCTATATCATGGCCGACAAGCAGAAGAACGGCATCAAGGTGAACTTCAAGATCCGCCACAACATCGAGGACGGCAGCGTGCAGCTCGCCGACCACTACCAGCAGAACACCCCCATCGGCGACGGCCCCGTGCTGCTGCCCGACAACCACTACCTGAGCACCCAGTCCAAGCTGAGCAAAGACCCCAACGAGAAGCGCGATCACATGGTCCTGCTGGAGTTCGTGACCGCCGCCGGGATCACTCTCGGCATGGACGAGCTGTACAAGTGAGGGCAGGCGAGCCCCACCCCGGCCCCGGCCCCTCCTGGACTCGCCTGCTCGCTTCCCCTTCCCAGGCCCGTGGCCAACCCAGCAGTCCTTCCCTCAGCTGCCTAGGAGGAAGGGACCCAGCTGGGTCTGGGCCACAAGGGAGGAGACTGC

where C terminal tagging with GFP is in green (codon-optimized) + short linker in purple. 150 bp homology arms (orange) were incorporated via PCR amplification from a synthesized (IDT), codon-optimized monomeric EGFP.

dsPCR product was purified and 150 ng was used directly for transfection together with gRNAs. 70-80% confluent 24-well plates of U2Os cells were transfected with 2 μg PEI (1 μg/ml), 150 ng of plasmid and 150 ng of the PCR product. In addition, hMEC-AP2-GFP were treated with 1 μM DNA-PKc inhibitor NU7441 for 48 h post transfection. Two days after transfection cells were treated with puromycin (1 μg /ml) to enrich for successfully transfected cells. After expansion, GFP-positive cells were sorted by FACS, and single clones were expanded and genotyped.

#### Generating the U2Os-AP2-halo and HeLa-AP2-halo cell lines

U2Os-AP2-halo and HeLa-AP2-halo cell lines were generated with the same protocol as the U2Os-AP2-GFP cell line. The donor template sequence was:

GGCCAGCATCCTGGGGGGCCTCGTCTCACCCCAGGGTCTCCCCTCACACAGGTTTACACGGTCGTGGACGAGATGTTCCTGGCTGG CGAAATCCGAGAGACCAGCCAGACGAAGGTGCTGAAACAGCTGCTGATGCTACAGTCCCTGGAGGGAAGTGCATCTGGGAGCTCAGGCGCTAGTGGTTCAGC GAGCGGGGCAGAAATCGGTACTGGCTTTCCATTCGACCCCCATTATGTGGAAGTCCTGGGCGAGCGCATGCACTACGTCGATGTTGGTCCGCGCGATGGCACCC CTGTGCTGTTCCTGCACGGTAACCCGACCTCCTCCTACGTGTGGCGCAACATCATCCCGCATGTTGCACCGACCCATCGCTGCATTGCTCCAGACCTGATCGGTAT GGGCAAATCCGACAAACCAGACCTGGGTTATTTCTTCGACGACCACGTCCGCTTCATGGATGCCTTCATCGAAGCCCTGGGTCTGGAAGAGGTCGTCCTGGTCAT TCACGACTGGGGCTCCGCTCTGGGTTTCCACTGGGCCAAGCGCAATCCAGAGCGCGTCAAAGGTATTGCATTTATGGAGTTCATCCGCCCTATCCCGACCTGGGA CGAATGGCCAGAATTTGCCCGCGAGACCTTCCAGGCCTTCCGCACCACCGACGTCGGCCGCAAGCTGATCATCGATCAGAACGTTTTTATCGAGGGTACGCTGC CGATGGGTGTCGTCCGCCCGCTGACTGAAGTCGAGATGGACCATTACCGCGAGCCGTTCCTGAATCCTGTTGACCGCGAGCCACTGTGGCGCTTCCCAAACGA

GCTGCCAATCGCCGGTGAGCCAGCGAACATCGTCGCGCTGGTCGAAGAATACATGGACTGGCTGCACCAGTCCCCTGTCCCGAAGCTGCTGTTCTGGGGCACC CCAGGCGTTCTGATCCCACCGGCCGAAGCCGCTCGCCTGGCCAAAAGCCTGCCTAACTGCAAGGCTGTGGACATCGGCCCGGGTCTGAATCTGCTGCAAGAA GACAACCCGGACCTGATCGGCAGCGAGATCGCGCGCTGGCTGTCGACGCTCGAGATTTCCGGCTGAGGGCAGGCGAGCCCCACCCCGGCCCCGGCCCCTC CTGGACTCGCCTGCTCGCTTCCCCTTCCCAGGCCCGTGGCCAACCCAGCAGTCCTTCCCTCAGCTGCCTAGGAGGAAGGGACCCAGCTGGGTCTGGGCCACAA GGGAGGAGACTGC

where C terminal tagging with halo is in green (codon-optimized) and short linker in purple. 150 bp homology arms (orange) were incorporated via PCR amplification from a synthesized (IDT), codon-optimized monomeric halo tag.

#### Generating the HeLa-AP2-GFP, MCF7-AP2-halo, HDF-AP2-halo and CAco2-AP2-halo cell lines

The most effective gRNA (TGCTACAGTCCCTGGAGTGA) was ordered as sgRNA from Synthego and Cas9 protein (purified in the lab) was used instead of plasmid. The donor templates for these cell lines to insert either EGFP or Halo tag were same as above, respectively.

2.10^5^ cells were combined with 4 μg Cas9 protein, 45 pmol sgRNA and 300 ng purified PCR product into a nucleofection cuvette. The nucleofection protocols were the following: MCF7-AP2-halo: EN130, buffer SF; HDF-AP2-halo: EO114, buffer SF; Caco2-AP2-halo: DG113, buffer SF; HeLa-AP2-GFP: CN114, buffer SF.

Cell lines were then treated with 1 μM DNA-PKc inhibitor NU7441 for 48 hours post nucleofection.

#### Generating the U2Os-AP2-GFP-ITGB5-mScarlet cell line

This cell line was produced by the same protocol as the U2Os-AP2-GFP cells with the following changes: The gRNA sequence was CAAATCCTACAATGGCACTG, and the donor template was:

GGTTTGAGTGTGTGAGCTAACATGTGTCCTCATCCTCTTCCCCGCCGTGTTCTGTAGGCTTCAAATCCATTATACAGAAAGCCTATCTCCACGCACACTGTGGACTTCA CCTTCAACAAGTTCAACAAATCATATAACGGCACTGTTGACGGAAGTGCATCTGGGAGCTCAGGCGCTAGTGGTTCAGCGAGCGGGGTGAGCAAGGGCGAGGC AGTGATCAAGGAGTTCATGCGGTTCAAGGTGCACATGGAGGGCTCCATGAACGGCCACGAGTTCGAGATCGAGGGCGAGGGCGAGGGCCGCCCCTACGAGG GCACCCAGACCGCCAAGCTGAAGGTGACCAAGGGTGGCCCCCTGCCCTTCTCCTGGGACATCCTGTCCCCTCAGTTCATGTACGGCTCCAGGGCCTTCATCAAG CACCCCGCCGACATCCCCGACTACTATAAGCAGTCCTTCCCCGAGGGCTTCAAGTGGGAGCGCGTGATGAACTTCGAGGACGGCGGCGCCGTGACCGTGACC CAGGACACCTCCCTGGAGGACGGCACCCTGATCTACAAGGTGAAGCTCCGCGGCACCAACTTCCCTCCTGACGGCCCCGTAATGCAGAAGAAGACAATGGGC TGGGAAGCATCCACCGAGCGGTTGTACCCCGAGGACGGCGTGCTGAAGGGCGACATTAAGATGGCCCTGCGCCTGAAGGACGGCGGCCGCTACCTGGCGG ACTTCAAGACCACCTACAAGGCCAAGAAGCCCGTGCAGATGCCCGGCGCCTACAACGTCGACCGCAAGTTGGACATCACCTCCCACAACGAGGACTACACCG TGGTGGAACAGTACGAACGCTCCGAGGGCCGCCACTCCACCGGCGGCATGGACGAGCTGTACAAGTAATGTTTCCTTCTCCGAGGGGCTGGAGCGGGGATCT GATGAAAAGGTCAGACTGAAACGCCTTGCACGGCTGCTCGGCTTGATCACAGCTCCCTAGGTAGGCACCACAGAGAAGACCTTCTAGTGAGCCTGGGCCAGGA GCCCACAGTGCCT

where A=silent mutations in 5’ HA, linker region is in purple, and mScarlet is in orange.

### Lentiviral shRNA production and transduction

Lentiviruses for shRNA production were produced using packaging plasmids pCMVR and pMD2.g and specific shRNAs in pLKO.1 vector as follows: 80% confluent HEK293T cells in DMEM supplemented with 10% FBS and 100U penicillin-streptomycin were transfected using PEI MAX transfection reagent. 5 hours later, the medium was changed to DMEM supplemented with 4% FBS and 25mM HEPES. Media containing lentiviral particles were harvested after 48 and 72 hours, filtered (0.45 μm), aliquoted and stored at -80°C.

U2Os-AP2-GFP cells were transduced with lentiviral media expressing respective shRNAs in the presence of Polybren 8 μg/ml (Sigma-Aldrich, TR-1003) for 5 hours, and replaced with culture medium. 48 h later, puromycin (1 μg/ml) was added for 24 h to allow selection of transduced cells. Experiments targeting AP2 subunits were performed otherwise similarly but without puromycin selection. All shRNA-silenced cell lines were replated on non-coated glass-bottomed imaging dishes one day prior to imaging.

The following sequences were targeted:

Integrin β5 targeting (ITGB5) shRNAs: TRCN0000057744 (GCATCCAACCAGATGGACTAT), TRCN0000057745 (GCTGTGCTATGTTTCTACAAA) Integrin β1 targeting (ITGB1) shRNAs: TRCN0000029644 (CCTGTTTACAAGGAGCTGAAA), TRCN0000029645 (GCCTTGCATTACTGCTGATAT)

AP2 sigma 1 targeting (AP2S1) shRNAs: TRCN0000060263 (GACGCCAAACACACCAACTTT), TRCN0000060266 (GTGGAGGTCTTAAACGAATAT), TRCN0000060267 (CACAACTTCGTGGAGGTCTTA).

AP2 alpha 1 (AP2A1) targeting shRNAs: TRCN0000065108 (GCTGAATAAGTTTGTGTGTAA), TRCN0000065109 (GCACATTGACACCGTCATCAA)

Integrin alpha 5 (ITGA5) targeting shRNAS: TRCN0000029651 (CCACTGTGGATCATCATCCTA), TRCN0000029652 (CCTCAGGAACGAGTCAGAATT).

Tensin1 targeting (TNS1) shRNAs: TRCN0000002953 (GAGGATAAGATTGTGCCCATT), TRCN0000002954 (CCCAAAGAAGGTACGTGCATT). Talin1 targeting (TLN1) shRNA: TRCN0000123106 (GCCTCAGATAATCTGGTGAAA)

### Clathrin exon 31 analysis

Cell culture dishes were coated with 10 μg/ml of FN in a cell culture incubator, or left uncoated. Next day, U2Os cells were plated to reach confluency in 24 h. RNA extraction, cDNA synthesis, and PCR followed those described by Moulay et al. (2020) with minor variations.

Total RNA was extracted from cells using TRIzol reagent with an additional acidic phenol (pH 5.4) extraction step to remove genomic DNA contamination. cDNA synthesis from 1 μg total RNA was carried out using Maxima H Minus Reverse Transcriptase (Thermo Fisher Scientific) and oligo dT12–18 (Life Technologies). No enzyme reactions were included to confirm that no genomic DNA was present. PCR was performed using Phusion High-Fidelity DNA polymerase (Thermo Fisher Scientific) with no other variations from Moulay et al., (2020). Primers used were: F’ TGC CCT ATT TCA TCC AGG TCA, R’ ATG GGT TGT GTC TCT GTA GC. Gel images were acquired using a GelDoc XL (Bio Rad).

### Western blots

U2Os were silenced for respective proteins as mentioned above, and lysed into 150 mM NaCl, 50 mM Tris pH8 and 1% Igepal (Sigma). Protein concentrations were measured with the Pierce BCA kit (Thermo Scientific), and boiled in Laemmli loading buffer with 10% mercaptoethanol. Equal amounts of protein lysates were loaded onto SurePAGE Bis-Tris 4-20% gels (Genscript) and transferred either to

0.2 μm PVDF or 0.45 μm nitrocellulose (GE Healthcare) membranes. Membranes were blocked in 5% BSA or skimmed milk for 1 h, incubated with primary antibodies at +4°C overnight, and HRP-conjugated secondary antibodies (1706516, 1721019, Biorad) for 1 h in RT. HRP was activated with Supersignal West Femto or Pico reagents (Thermo Scientific) and bands were detected with ChemiDoc XRS+ (BioRad).

### Microscopy

All live videos and images from fixed samples were acquired with the ONI Nanoimager microscope equipped with 405, 488, 561 and 647 lasers, an Olympus 1.49NA 100x super achromatic objective and a Hamamatsu sCMOS Orca flash 4 V3 camera.

#### Live time-lapse TIRF imaging Endogenous AP2 life-time monitoring

35 000 U2Os-AP2-GFP cells were plated on precoated/non-coated areas of the dish resulting in approximately 70-80% confluency 16-20 h later, at the onset of imaging. Alternatively, shRNA-silenced cell lines were plated. After overnight culture, 25 μM HEPES was added and samples were subjected to live TIRF imaging in a pre-heated +37°C chamber.

The ONI nanoimager microscope set to TIRF angle was used to acquire AP2 lifetimes at the cell membrane from 300 frames (1 frame/s) with the exposure time of 330 ms. Each video represents endocytic events from 2-3 cells (total field of view).

#### Acute manipulation of integrin activity

##### Integrin β1 blocking

Acute modulation of ligand binding activity for integrin β1 was achieved using the function blocking antibody mab13 (0.3 μg/ml). U2Os-AP2-GFP cells were plated on FN as explained above, and 16-20 h later subjected to live TIRF imaging. 0 min sample has no mab13 added, to control base-line FCL proportions. Immediately after mab13 addition, 5 min time lapses were continuously collected until 35 min, control videos (time point 0) had no mab13 added.

##### Integrin β5 blocking

To acutely induce the inhibition of integrin αvβ5 we used the small molecular inhibitor Cilengitide (MedChem Express HY-16141, 10 μM). U2Os-AP2-GFP cells plated on non-coated imaging dishes were treated with Cilengitide for 15 or 45 min, fixed, stained, and imaged with the ONI nanoimager microscope at TIRF angle, and analyzed for the resulting reduction of RA coverage.

##### Cilengitide washout

U2Os-AP2-GFP-ITGB5-mScarlet cells were plated on non-coated imaging dishes and 1 d later confluent monolayers were treated with 1 μM Cilengitide for 15-25 min, during which most FCLs and RAs were dissociated from the cell membrane. Samples were then washed twice and immediately subjected to live TIRF imaging to detect the de novo formation of FCLs and RAs. 1 h time-lapses were acquired with the ONI nanoimager microscope at TIRF angle, at 30 s intervals, with an exposure time of 100 ms for AP2 and 300 ms for integrin β5.

### Immunofluorescence experiments

#### Immunofluorescent staining and imaging

For immunofluorescence experiments, cells were fixed with 4% paraformaldehyde-PBS for 15 min in a +37°C incubator, washed with PBS and blocked with 1% BSA-PBS. Primary antibodies diluted in 1% BSA-PBS were incubated for 1 h, samples were washed with PBS, and secondary antibodies diluted in 1% BSA-PBS were let to bind for 30 min. Samples were imaged with the ONI nanoimager microscope using TIRF angle and exposure times of 500 ms or 1000 ms.

#### Manipulation of CME machinery to study RA formation

Clathrin assembly at the cell membrane was reduced by silencing two subdomains of AP2, or by overexpressing AP180ct, which acts as a dominant negative for AP2. U2Os AP2-GFP cells silenced for AP2A1 shRNA #1 and #2 or AP2S1 shRNA #1, #2 and #3, and control shRNA, and plated on non-coated imaging dishes were fixed and stained for integrin αvβ5 and FA marker p-Pax. Alternatively, AP180ct was overexpressed in U2Os AP2-GFP cells and cells were plated on non-coated imaging dishes, fixed the next day, and stained for integrin αvβ5 and p-Pax. RA formation (integrin αvβ5 adhesions w/o FA marker) and FAs (integrin αvβ5 colocalising with FA marker) were imaged using TIRF microscopy.

#### Manipulation of integrin activity and availability

We performed time-series experiments to block integrin β1 active conformation with the mab13 antibody or to inhibit integrin αvβ5 with Cilengitide. Experiments utilizing mab13 were performed with U2Os-AP2-GFP cells plated on FN-coated dishes to disfavour the preformation of FCLs and RAs. Mab13 (0.3 μg/ml) was added to replicate samples and fixed 15, 30 or 45 min later. Similarly, U2Os-AP2-GFP cells plated on non-coated dishes to favour the preformation of FCLs and RAs were treated with Cilengitide. Samples were stained for integrin αvβ5 and FA marker p-Pax, and imaged using TIRF microscopy.

#### Cilengitide washout

Cilengitide washout experiments were developed to study acute reappearance of FCLs and RAs in cell cultures. Cells were treated for 15-25 min with 1 μM Cilengitide (or DMSO as control) and washed two times. Unwashed controls were collected. Washed samples were let to recover for indicated time points, fixed, stained for p-Pax, subjected to TIRF imaging, and analysed for the reappearance of FCLs and RAs (RA coverage).

U2Os-AP2-GFP cells silenced for integrin β1, integrin β5, integrin α5 or Tensin1 and controls were plated either on FN coated dishes (control shRNA and ITGB1 shRNAs, ITGA5 shRNAs or TNS1 shRNAs) or on non-coated dishes (control shRNA and ITGB5 shRNAs), and fixed the next day, 16-20 h later. Samples were stained for integrin αvβ5 and FA marker p-Pax and imaged using TIRF microscopy.

#### Micropatterning

U2Os-AP2-GFP or U2Os-AP2-GFP-ITGB5-mScarlet cells were seeded onto micropatterned glass coverslips (CYTOO) either precoated with FN 10 μg/ml or left uncoated. Excess amount (60 000) of cells were plated, and monitored for attachment to the patterns. The cells plated on FN had attached in 1 h, and the cells plated on non-coated micropatterns had attached in 4 h. After attachment, excess cells were carefully rinsed, and the samples were fixed 16-20 h later, stained, and subjected to immunofluorescence analysis with TIRF microscopy.

#### Wound healing-stimulated migration

U2Os-AP2-GFP-ITGB5-mScarlet cells were plated confluent onto non-coated imaging dishes for 2 days. Fully confluent monolayers were wounded with a micropipette tip, washed twice with fresh complete medium and let to migrate. 0 min samples were collected directly after wounding. After fixing, samples were stained and subjected to TIRF imaging. Tile images (5×1), starting at the edge of the wound were acquired with a 25% overlap and stitched using the pairwise stitching plug-in Image J.

### Image analyses

#### CME lifetime analyses (FCL proportion)

To track CME events and measure lifetimes we used ‘u-track 2.0’ multiple-particle tracking MATLAB software at default settings (Jaqaman et al., 2008). For all experiments, “n” refers to a movie, which contained 2-4 cells. To determine the proportion of FCLs, we used the output from u-track to count the number of pits (events lasting longer than 20 s and shorter than 120 s) and the number of FCLs (events lasting longer than 120 seconds, as described in (Saffarian et al., 2009) in all frames. We took a conservative approach to identify CCPs, where events that were present at the start or lasted beyond the end of the movies were not counted as CCPs. This approach artificially led to higher FCL proportions in the first and final 120 movie frames. Therefore, FCL proportions are presented as the average FCL proportion from frames 120 to 175 for each movie.

#### Other analyses

With the exception of FCL proportions, all image analyses were performed using ImageJ. Simple fluorescence measurements were done manually. Others were performed using custom scripts as shown below:

##### RA coverage

Individual cells were marked and ITGB5 (or αvβ5) and p-Pax channels were segmented using the Robust Automatic Threshold Selection function. RAs were defined as ITGB5 (or αvβ5) signals not colocalizing with p-Pax. The area of RAs was then divided by the area of each marked cell to obtain RA coverage. Data is presented as percentage of the cell area covering RAs.

For the wound healing experiment, a line on the migration front of each image was manually drawn. This line was then used as a reference to automatically draw a box, 100 pixels in width (11.7 μm). RA coverage (as above) was calculated for this box, which was then moved inward in the culture in 50 pixels steps, where the RA coverage analysis was repeated. Values are normalized to the average RA coverage on the three innermost areas in the culture.

##### AP2-ITGB5 dynamics

Events showing the appearance of both AP2 and ITGB5 were identified by visual inspection of videos. For the generation of graphs, we selected only events where we could unambiguously ensure that significant FCLs and ITGB5 signals were not present in the region for at least 3 minutes. Time zero was defined as the frame where AP2 signal appeared and fluorescence Intensity from a 10 μm x 10 μm region around each event was measured for 3 min before (six frames) and 5 min after (10 frames). Fluorescence was normalised to the highest value in these frames.

##### AP2 intensity per colocalisation status

AP2, ITGB5 and p-Pax channels were segmented using the Robust Automatic Threshold Selection function. Each segmented AP2 spot had the fluorescence intensity measured from the original image and classified for its colocalisation with either marker (ITGB5 or p-Pax). We used full images for these analyses.

##### RAs colocalising to AP2

Individual cells were marked and AP2, ITGB5 and p-Pax channels were segmented using the Robust Automatic Threshold Selection function. RAs were defined as ITGB5 signals not colocalizing with p-Pax. In the conditions used for these experiments, (FN + mab13) RAs were primarily individual spots. The colocalisation of RAs to AP2 was classified by measuring the intensity of each RA region at the segmented AP2 channel.

##### FN intensity vs. AP2 intensity

AP2, ITGB5 and P-PAX channels were segmented using the Robust Automatic Threshold Selection function. Each segmented AP2 spot had the fluorescence intensity measured from the original image. A 3 μm x 3 μm region was drawn around each AP2 spot and used to measure the intensity of FN from the original image. Data is presented as the fluorescence for each AP2 spot.

### Statistics

Figure legends state the exact n-values and individual repeats used in analyses. For multiple comparisons one-way ANOVA was performed followed by Tukey’s multiple comparison. Pairwise comparisons were performed using two-tailed Student’s t-test with equal variance. All graphs and statistical calculations were performed with GraphPad Prism 9.

