## Supplementary figures 1-5 for "Reticular Adhesion Formation is Mediated by Flat Clathrin Lattices and Opposed by Fibrillar Adhesions"

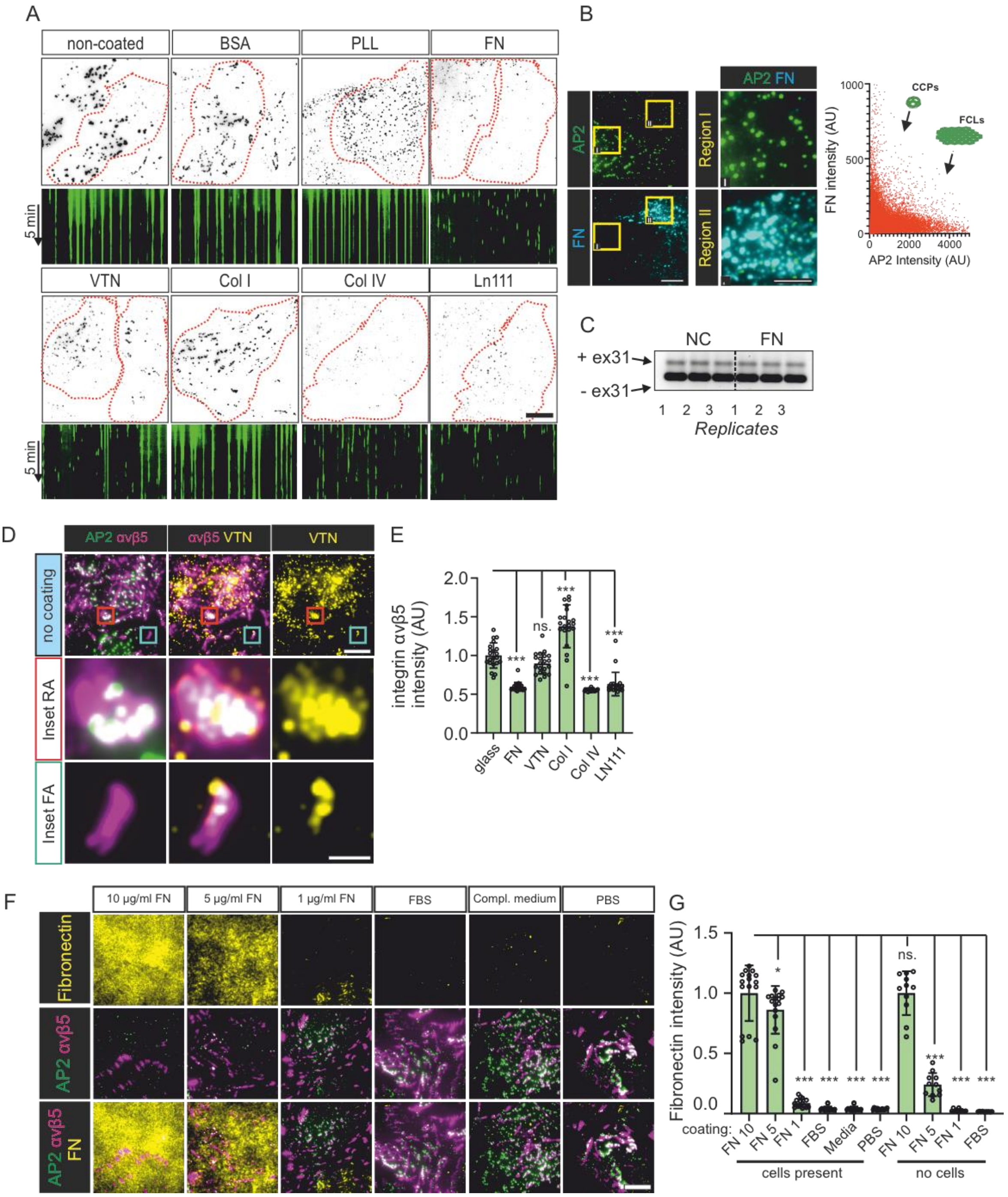


**Figure S1 - (A)** U2Os-AP2-GFP cells plated on PLL, BSA, FN, VTN, Col I, Col IV, LN111-coated or non-coated dishes overnight. Samples were imaged with TIRF microscopy at 1 s intervals for 5 min. Representative 15 s time projections, and 5 min kymographs of time-lapse videos from samples in Fig 1B. **(B)** Left: U2Os-AP2-GFP cells plated on non-coated dishes overnight were stained for fibronectin (FN). Representative TIRF images. Right: graph showing the reduced brightness of AP2 in regions with higher FN signal (measured from a 1.5 µm x 1.5 µm region around each AP2 spot, n=32988 AP2 spots, 27 images from one representative sample). **(C)** U2OS cells plated on non-coated (NC) or FN-coated (FN 10 µg/ml) dishes were analyzed for clathrin exon 31 density by RT-PCR, n=3 biological replicates. **(D)** U2Os-AP2-GFP were plated to non-coated dishes and stained for integrin αvβ5 and vitronectin (VTN). Representative TIRF images. **(E)** Analysis of integrin αvβ5 fluorescent intensity of samples from Fig 2A. N (images):

FN=15, VTN/Col I/LN111/non coated=21, Col IV=17. Results were obtained from one representative experiment, similar results were observed in four individual experiments. $F(5, 120)=85.49$, $P<0.0001$. **(F)** U2Os-AP2-GFP cells plated on 10 µg/ml FN, 5 µg/ml FN, 1 µg/ml FN, FBS or complete MEM medium-coated dishes were stained for fibronectin and integrin αvβ5. Representative TIRF images. **(G)** Fibronectin integrated fluorescent density of dishes coated as in G, and plated or not plated with U2Os cells. N=16-10/sample, from two independent experiments. $F(9, 129)=184.8$, $P<0.0001$. Data are the mean ± SD, ns. non-significant p-value; *** p-value < 0.001. Scalebars 10 µm and 5 µm insets except in D insets are 2 mm.

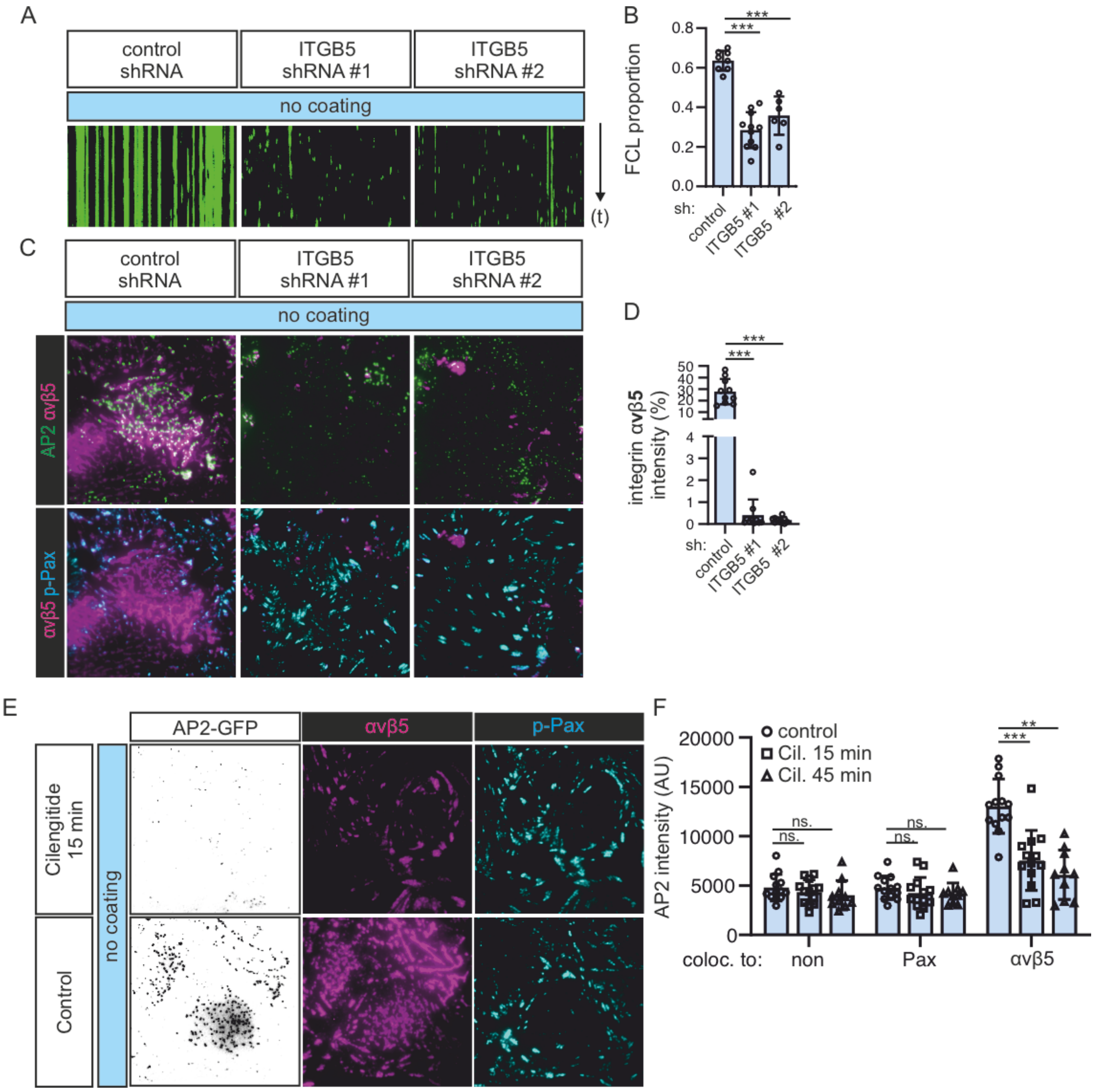


**Figure S2. (A)** U2Os-AP2-GFP cells silenced for integrin β5 (ITGB5) with two shRNAs (shITGB5 #1, #2) or control, were imaged using TIRF microscopy at 1 s intervals for 5 min. Representative 5 min kymographs. **(B)** Analysis of FCL proportions from time-lapse videos in A. N (videos): control=8, shITGB5 #1=11, shITGB5 #2=6, from three independent experiments. One-way ANOVA with Tukey's multiple comparison, $F(2, 22)= 44.46$, $P<0.0001$. **(C)** U2Os-AP2-GFP cells silenced for integrin β5 (ITGB5) with two shRNAs (shITGB5 #1, #2) or control, were stained for integrin αvβ5 and p-Pax. Representative TIRF images. **(D)** Analysis of integrin β5 silencing efficiency, n=10, from one representative experiment. Similar results were observed from three independent experiments. One-way ANOVA with Tukey's multiple comparison, $F(2, 27)= 63.29$, $P<0.0001$. **(E)** U2Os-AP2-GFP cells plated on non-coated dishes for 20 h were treated with the integrin αvβ5 inhibitor Cilengitide (10 µM) for 15 or 45 min and stained for integrin αvβ5 and p-Pax. Representative TIRF images. **(F)** Analysis of AP2 signal colocalising with p-Pax or integrin αvβ5 over time from samples in E. N (images): control=14, Cil 15min=13, Cil 45 min=10, from one representative experiment. Similar results were observed in four independent experiments. One-way ANOVA and Tukey's multiple comparison, $F(2, 99)=63.38$, $P<0.0001$**.** Data are the mean ± SD, ns. non-significant p-value; *** p-value < 0.001. Scalebars 10 µm.

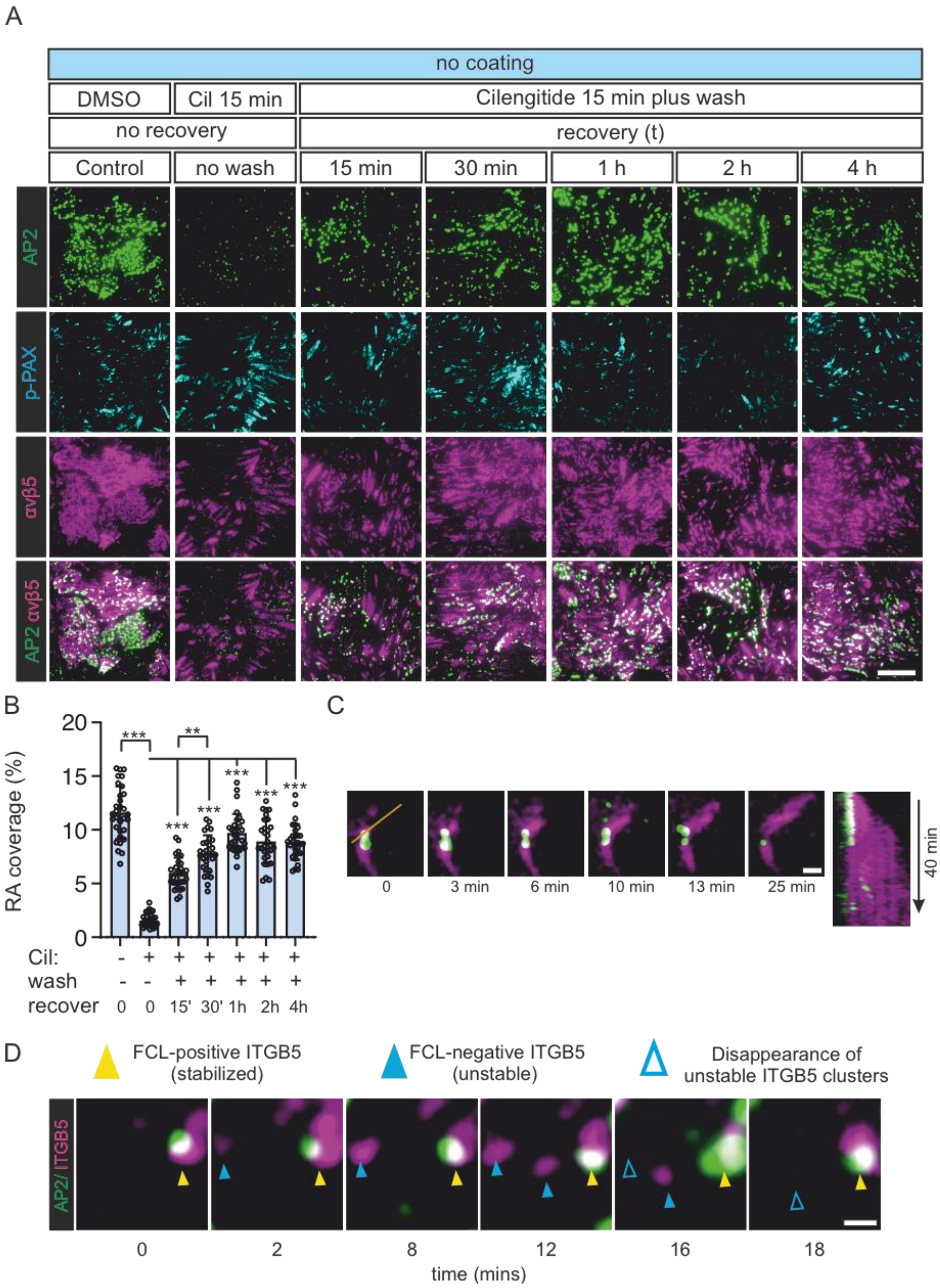


**Figure S3. (A)** U2Os-AP2-ITGB5-mScarlet cells plated on non-coated dishes were treated with Cilengitide (10 μM) or DMSO for 25 minutes. Samples given Cilengitide were washed twice with fresh complete medium (except no wash-control), fixed at the respective recovery time points and stained for p-Pax. Representative TIRF images. **(B)** Analysis of RA coverage for samples in A. N (images): control=31, no wash=32, 15 min recovery=30, 30 min recovery=30, 1 h recovery=31, 2 h recovery=30, 4h recovery=30, from one representative experiment. Similar results were observed in three independent experiments. **(C-D)** Additional examples of individual ITGB5 and AP2 events from experiments shown in figure 5. In C, an event where an RA grows from a stabilized FCL/ITGB5 cluster. A kymograph for the line on time 0 is shown on the right. In D, 3 events are shown. One FCL-stabilized ITGB5 cluster (yellow arrows) and two non-stabilized ITGB5 clusters (blue arrows). Open blue arrows represent frames post disappearance of ITGB5 clusters. Scalebars 10 μm.

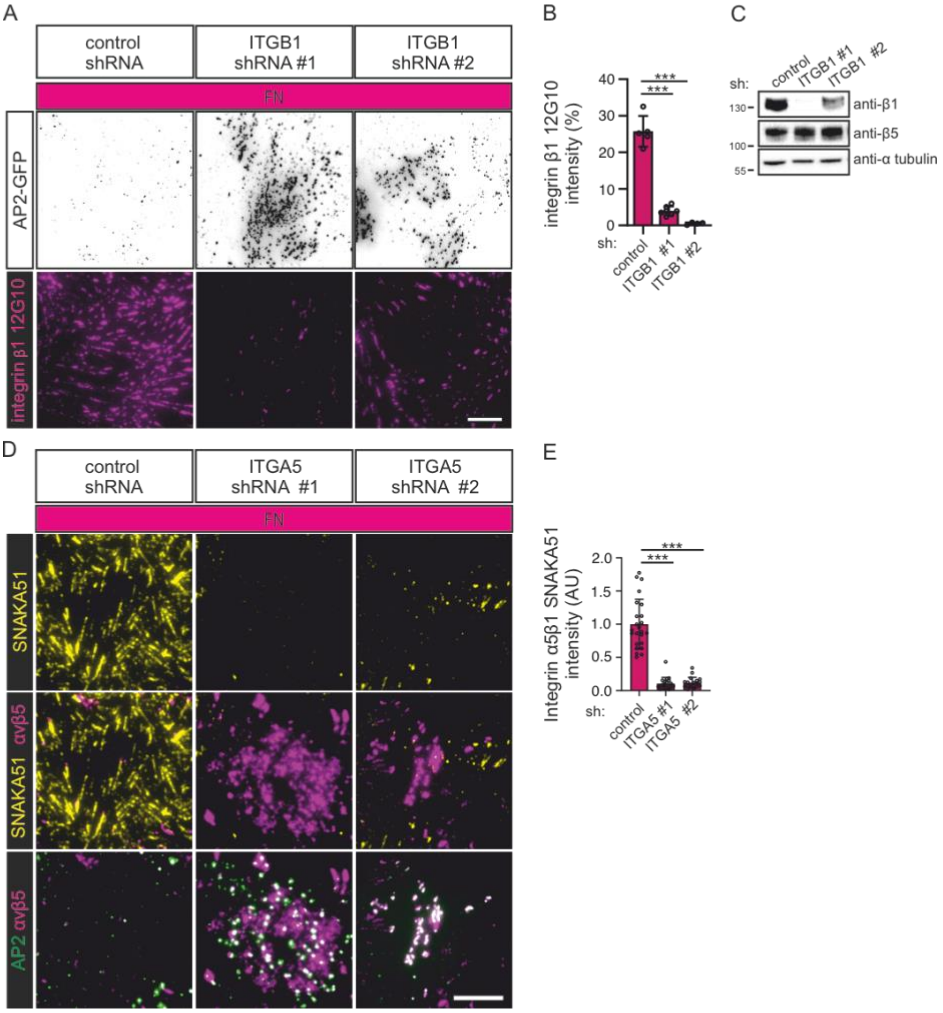


**Figure S4. (A)** U2Os-AP2-GFP cells silenced for integrin β1 with two different shRNAs (shITGB1 #1, #2) or control shRNA, were plated on FN-coated dishes and stained for active integrin β1 (12G10 antibody). Representative TIRF images. **(B)** Analysis of 12G10 fluorescent intensity. N=7 from one representative experiment. Similar results were observed from two individual experiments. One-way ANOVA with Tukey's multiple comparison, $F(2, 13)=149.9$, $P<0.0001$. **(C)** Western blots showing integrin β1 silencing efficiency and the effect on integrin β5 protein levels. U2Os cells silenced for integrin β1 shRNAs (shITGB1 #1, #2) or control shRNA, were blotted for integrin β1, integrin α5, and α-tubulin. Representative blots out of two individual experiments. **(D)** U2Os-AP2-GFP-ITGB5-mScarlet cells silenced for integrin α5 with two different shRNAs (shITGA5 #1, #2) or control shRNA were plated on FN-coated dishes and stained for integrin α5 (SNAKA51 antibody). Representative TIRF images. **(E)** Analysis of SNAKA51 fluorescent intensity from widefield microscopic images. N (images): shScr=24, shITGA5 #1=20, shITGA5 #2=19, from one representative experiment. Similar results were observed in two individual experiments. One-way ANOVA with Tukey's multiple comparison, $F(2, 60)=1$, $P<0.0001$. Data are the mean ± SD,*** p-value < 0.001. Scalebars 10 µm.

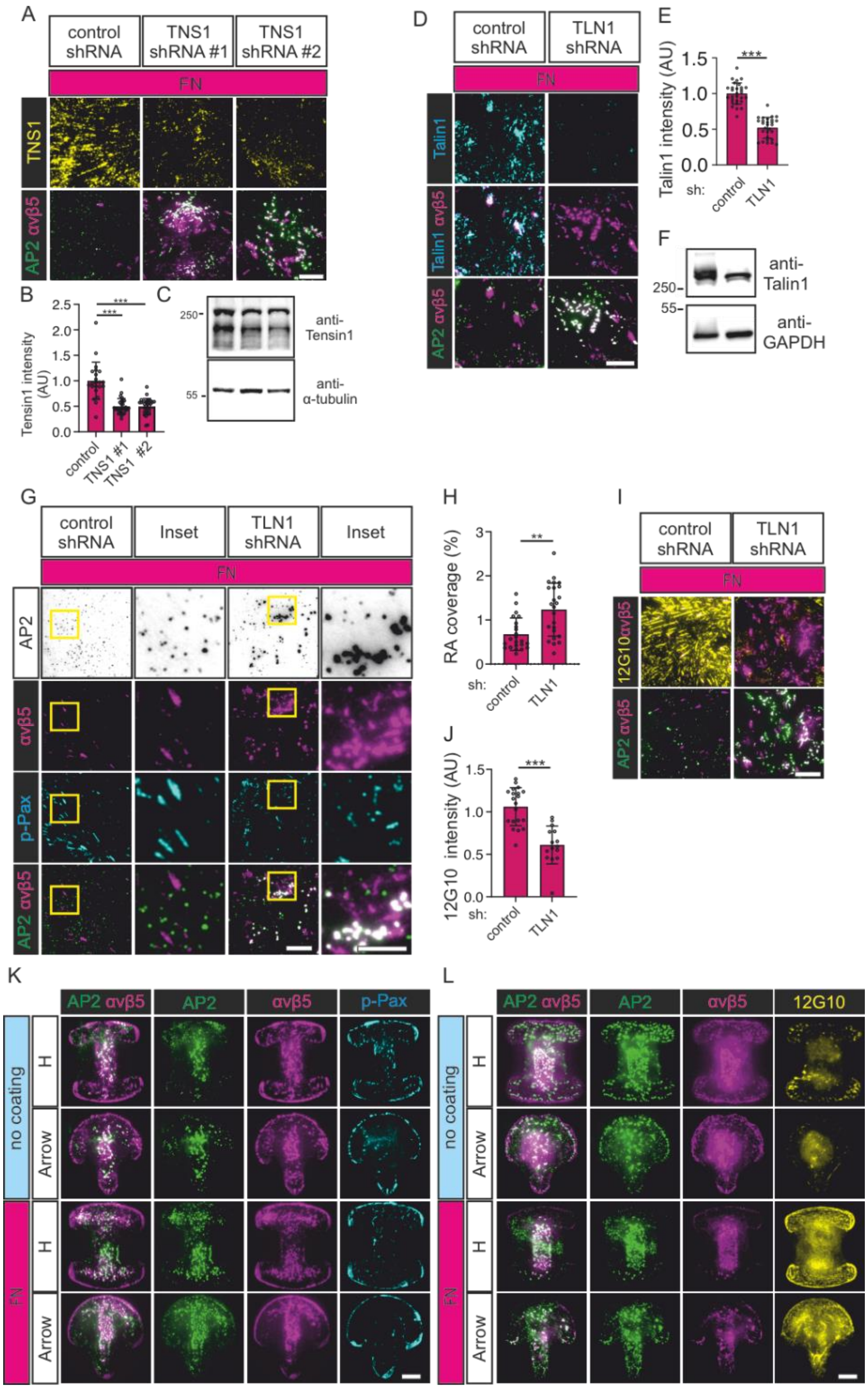


**Figure S5. (A)** U2Os-AP2-GFP-ITGB5-mScarlet cells silenced for Tensin1 with two different shRNAs (shTNS1 #1, #2) or control shRNA, were plated on FN-coated dishes and stained for Tensin1. Representative TIRF images. **(B)** Analysis of Tensin1 fluorescent intensity from samples in A. N (images): shScr control=22, shTNS1 #1=29, shTNS1 #2=31, from one representative experiment. Similar results were observed in three individual experiments. One-way ANOVA with Tukey's multiple comparison, $F(2,\ 79)=38.09$, $P<0.0001$. **(C)** Representative western blots of Tensin1 silencing efficiency. U2Os-AP2-GFP-ITGB5-mScarlet cells silenced for Tensin1 with two different shRNAs (shTNS1 #1, #2) or control shRNA were blotted for Tensin1 and α-tubulin. **(D)** U2Os-AP2-GFP-ITGB5-mScarlet cells silenced for Talin1 with shRNA or control shRNA were plated on FN-coated dishes and stained for Talin1. Representative TIRF images. **(E)** Analysis of Talin1 fluorescent intensity from samples in D. N (images): shScr=32, shTLN1=31, from two individual experiments. Two-tailed Student's t-test, P<0.0001. **(F)** Representative western blots of Talin1 silencing efficiency. U2Os-AP2-GFP-ITGB5-mScarlet cells silenced for Talin1 shRNA or control shRNA were blotted for Talin1 and GAPDH. **(G)** U2Os-AP2-GFP-ITGB5-mScarlet cells silenced for Talin1 with shRNA or control shRNA were plated on FN coated dishes and stained for p-Pax. Representative TIRF images. **(H)** Analysis of RA coverage from samples in G. N (images): n=23. Two-tailed Student's t-test, P<0.0001. **(I)** U2Os-AP2-GFP-ITGB5-mScarlet cells silenced for Talin1 with shRNA or control shRNA were plated on FN-coated dishes and stained for integrin β1 12G10. Representative TIRF images. **(J)** Analysis of 12G10 fluorescent intensity with samples from I. N (images): shScr=20, shTLN1=15, from one representative experiment. Similar results were observed in two individual experiments. Two-tailed Student's t-test, P<0.0004. **(K)** U2Os-AP2-GFP cells were grown on FN-coated or non-coated micropatterns (1100 mm2) and stained for integrin αvβ5 and p-Pax. Representative TIRF images. **(L)** U2Os-AP2-GFP-ITGB5-mScarlet cells were grown on FN-coated or non-coated micropatterns (1100 mm2) and stained for active integrin β1 12G10. Representative TIRF images. Data are the mean ± SD,** p-value < 0.01, *** p-value < 0.001. Scalebars 10 µm, insets 5 µm.
